# Septins function in exocytosis via physical interactions with the exocyst complex in fission yeast cytokinesis

**DOI:** 10.1101/2024.07.09.602728

**Authors:** Davinder Singh, Yajun Liu, Yi-Hua Zhu, Sha Zhang, Shelby M. Naegele, Jian-Qiu Wu

## Abstract

Septins can function as scaffolds for protein recruitment, membrane-bound diffusion barriers, or membrane curvature sensors. Septins are important for cytokinesis, but their exact roles are still obscure. In fission yeast, four septins (Spn1 to Spn4) accumulate at the rim of the division plane as rings. The octameric exocyst complex, which tethers exocytic vesicles to the plasma membrane, exhibits a similar localization and is essential for plasma membrane deposition during cytokinesis. Without septins, the exocyst spreads across the division plane but absent from the rim during septum formation. These results suggest that septins and the exocyst physically interact for proper localization and function. Indeed, we predicted six pairs of interactions between septin and exocyst subunits by AlphaFold, most of them are confirmed by co-immunoprecipitation and yeast two-hybrid assays. Exocyst mislocalization results in mistargeting of secretory vesicles and their cargos, which leads to cell-separation delay in septin mutants. Our results indicate that septins guide the targeting of exocyst complex on the plasma membrane for vesicle tethering during cytokinesis through physical interactions.

## Introduction

Septins are a family of GTP-binding proteins that are highly conserved from yeast to mammalian cells (Longtine et al., 1996; Gladfelter et al., 2001; Cao et al., 2007; Nishihama et al., 2011; Onishi and Pringle, 2016; Marquardt et al., 2019; Woods and Gladfelter, 2021). They form hetero-oligomeric complexes that can assemble into different higher-order structures such as rings, gauzes, hourglasses, and carry out various functions (Frazier et al., 1998; Hsu et al., 1998; Kinoshita, 2003; Sheffield et al., 2003; Bertin et al., 2008; Garcia et al., 2011; Bridges et al., 2014). Septins can serve as scaffolds for protein recruitment at discrete cellular locations (Gladfelter et al., 2001; Versele and Thorner, 2005; Mostowy and Cossart, 2012; Finnigan et al., 2015; Meitinger and Palani, 2016; Perez et al., 2016; Marquardt et al., 2019). Septins are proposed to act as a diffusion barrier to ensure that cellular components are spatially segregated or compartmentalized (Dobbelaere and Barral, 2004; Caudron and Barral, 2009; Hu and Nelson, 2011; McMurray et al., 2011). They can also sense the membrane curvatures and/or deform the plasma membrane due to their lipid-binding properties (Bridges and Gladfelter, 2016; Cannon et al., 2017; Cannon et al., 2019; McMurray, 2019; Shi et al., 2023). The diverse roles of septins lead to their involvement in multiple processes including cytokinesis, mitosis, exocytosis, apoptosis, fungal or viral infections, neuronal spine morphogenesis, ciliogenesis, and spermiogenesis (Longtine et al., 1996; Kartmann and Roth, 2001; Mostowy and Cossart, 2012; Dolat et al., 2014; Momany and Talbot, 2017; Ageta-Ishihara and Kinoshita, 2021; Neubauer and Zieger, 2021; Woods and Gladfelter, 2021; Safavian et al., 2023).

One of the best-studied septin functions is their roles in cytokinesis in the budding yeast *Saccharomyces cerevisiae* (Hartwell, 1971; Haarer and Pringle, 1987; Ford and Pringle, 1991; Kim et al., 1991; DeMarini et al., 1997; Bi et al., 1998; Lippincott and Li, 1998; Longtine et al., 1998). The septin ring or hourglass structures at the presumptive bud site and bud neck are required for the recruitment and maintenance of various cytokinesis proteins (Bi et al., 1998; Lippincott and Li, 1998; Gladfelter et al., 2001; Longtine and Bi, 2003; Marquardt et al., 2019). These roles occur through either direct interactions with proteins such as the F-BAR protein Hof1 (Vallen et al., 2000; Meitinger et al., 2013; Oh et al., 2013), or through septin-binding proteins such as Bni5, which links septins to the myosin-II heavy chain Myo1 (Lee et al., 2002; Fang et al., 2010; Finnigan et al., 2015). During cytokinesis, the septin double rings were proposed to function as a diffusion barrier for proteins such as the exocyst component Sec3 and chitin synthase II Chs2 at the division site (Dobbelaere and Barral, 2004). But other studies have challenged this view by showing that Chs2 localizes efficiently to the division site in the absence of septin rings (Wloka et al., 2011). Regardless of the debate, septins are known to play essential roles in budding yeast cytokinesis. However, no physical interactions between septins and the exocyst have been reported, even in the genome wide interactome studies (Michaelis et al., 2023). Moreover, budding yeast septins also serve as scaffolds for the localization of hundreds of proteins at the bud neck including signaling proteins, bud site selection proteins, and chitin synthases (Longtine et al., 1996; DeMarini et al., 1997; Gladfelter et al., 2001; Longtine and Bi, 2003; Marquardt et al., 2019).

Unlike in budding yeast, septins are not essential in the fission yeast *Schizosaccharomyces pombe* and their roles in cytokinesis remain obscure (Longtine et al., 1996; Berlin et al., 2003; Tasto et al., 2003; Wu et al., 2010; Zheng et al., 2024). Fission yeast has seven septins, Spn1 to Spn7, with Spn1 to Spn4 expressing in vegetative cells and functioning at the division site (Longtine et al., 1996; Berlin et al., 2003; Tasto et al., 2003; An et al., 2004; Petit et al., 2005; Onishi et al., 2010; Wu et al., 2010). None of the septins Spn1-Spn4 is essential, but loss of some or all of them causes a delay in cell separation, resulting in multi-septated phenotype (An et al., 2004). Spn1 and Spn4 are the more important components of the septin ring structure during cytokinesis (An et al., 2004). Septins accumulate to the division site shortly before the contractile ring constriction and form a single ring which quickly transitions into unconstricting double rings (Berlin et al., 2003; Tasto et al., 2003; Wu et al., 2003; Wu et al., 2010). It is only known that septins recruit the anillin Mid2 (Berlin et al., 2003; Tasto et al., 2003), the guanine nucleotide exchange factor Gef3 (Muñoz et al., 2014; Wang et al., 2015), the small GTPase Rho4 (Wang et al., 2015), and the glucanases Eng1 and Agn1 to the division site (Martin-Cuadrado et al., 2005). Thus, we know much less about fission yeast septins compared to those in budding yeast. It remains mysterious what the most conserved functions of septins are during evolution.

Previous studies have suggested that septins function in exocytosis (Hsu et al., 1998; Vega and Hsu, 2003; Martin-Cuadrado et al., 2005; Perez et al., 2015; Tokhtaeva et al., 2015). Fission yeast septins are proposed to work with the exocyst complex to regulate the secretion of glucanases at the appropriate location, but it was reported that septins and the exocyst are independent for localization (Martin-Cuadrado et al., 2005; Perez et al., 2015). The exocyst is a highly conserved, octameric complex (Sec3, Sec5, Sec6, Sec8, Sec10, Sec15, Exo70, and Exo84) in exocytosis (TerBush et al., 1996; Wang et al., 2002; Hsu et al., 2004; Heider and Munson, 2012; Liu and Guo, 2012). It functions in late stages of exocytosis by promoting the tethering and fusion of post-Golgi secretory vesicles to the plasma membrane (TerBush et al., 1996; Hsu et al., 2004; Liu et al., 2018). Although studies suggest that septins regulate the exocyst complex and a possible involvement of septins in targeting secretory vesicles to the exocytic sites (Hsu et al., 1998; Vega and Hsu, 2003; Li et al., 2007; Gupta et al., 2015), no direct physical interactions between septins and the exocyst subunits have been reported in budding yeast or other organisms.

Here we report septins regulate the exocyst localization and vesicle targeting in fission yeast via physical interactions. We find that the loss of septin rings alters the exocyst localization, with increased concentration to the center and reduced localization to the rim of the division plane. The initial recruitment of the exocyst is independent of septins, but the exocyst requires septin rings to maintain the rim localization during furrow ingression. Consistently, we found multivalent physical interactions consistent with direct binding between septins and the exocyst subunits. Loss of the exocyst ring leads to abnormal accumulation of secretory vesicles in septin mutants. As a result, the glucan synthase Bgs1/Cps1 accumulates more to the center and the glucanase Eng1 is missing from the rim of the division plane, contributing to delayed cell separation and a thicker septum in septin mutant cells. Our findings provide insights into the regulation of the exocyst localization and function on the plasma membrane by septins in other systems.

## Results

### The septin and exocyst complex colocalize and are partially interdependent for localization at the division site

Both septins and the exocyst complex localize to the division site during cytokinesis (Longtine et al., 1996; Wang et al., 2002; An et al., 2004; Petit et al., 2005). To understand if and how they work together, we first examined the colocalization of the septin Spn1 and the exocyst subunit Sec3 in fission yeast. Spn1 is a key component in septin structures, and its deletion leads to a complete loss of all septins from the division site (An et al., 2004). Sec3 is a spatial landmark for exocytosis in budding yeast (Finger et al., 1998; Boyd et al., 2004; Luo et al., 2014). The fission yeast Sec3 is an essential gene and crucial for exocyst localization (Kim et al., 2010; Bendezu et al., 2012; Jourdain et al., 2012). Spn1 and Sec3 colocalized at the division site as a single ring first, and later as double rings during septum formation (Figure 1, A and B). The colocalization of Spn1 with another exocyst subunit Exo70 was confirmed using SoRa (Super resolution by Optical Re-Assignment) spinning disk confocal microscopy (Fig 1C). Sec3 and Exo70, but not Spn1, also concentrated at cell tips (Figure 1, C and D). However, Sec3 arrived at the division site 13.4 ± 2.2 min after spindle pole body (SPB) separation, about 10 min earlier than Spn1 that arrived at 23.6 ± 1.8 min (Figure 1, D and E). Time lapse movies of Exo70-tdTomato and Spn1-mEGFP confirmed that the exocyst appeared at the division site earlier than septins (Video 1). These observations suggest the spatial proximity between septins and the exocyst during certain stage of cytokinesis, raising the possibility of their functional coordination, which we would further investigate below.

**Figure 1.**
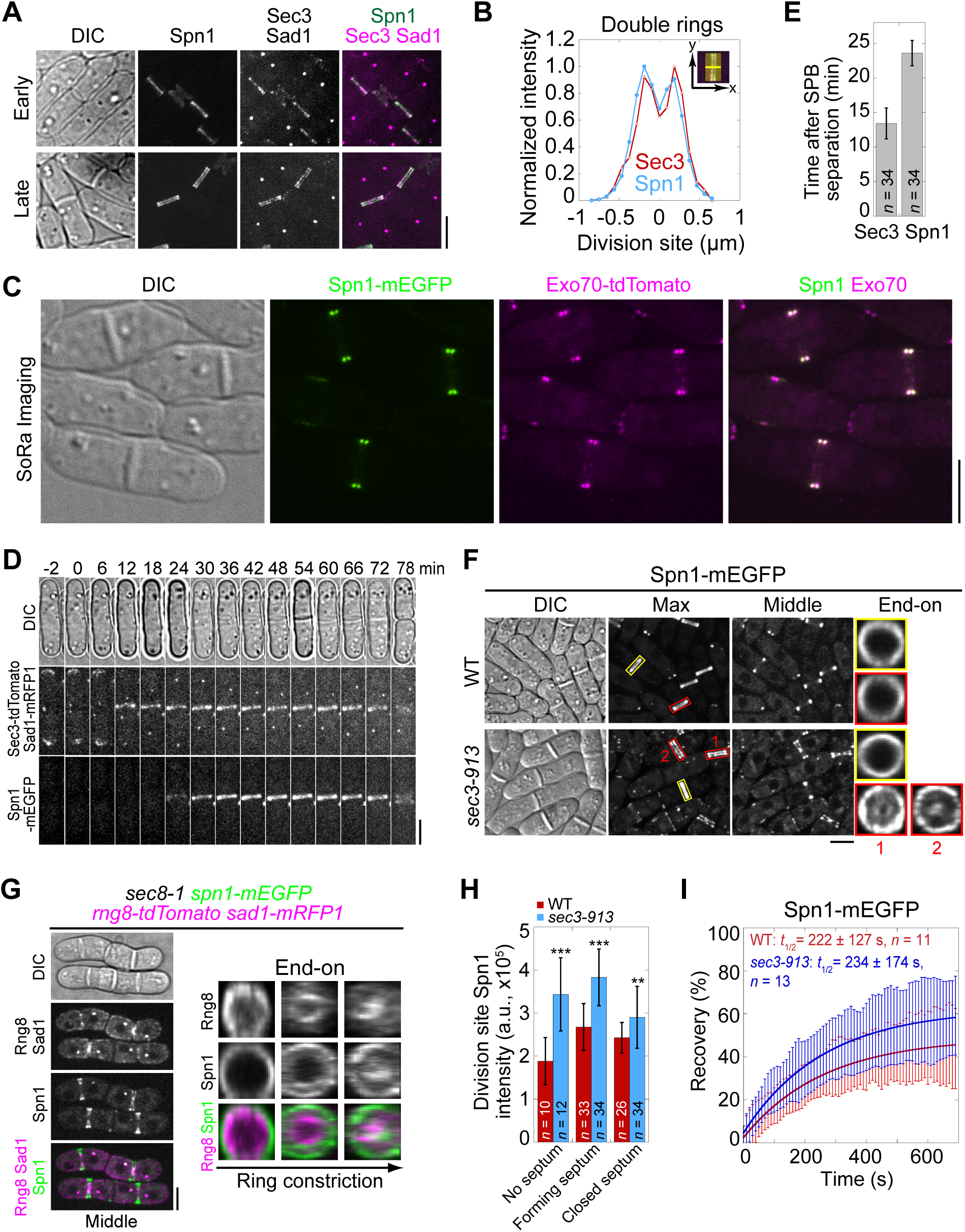
Septins and the exocyst colocalize at the division site and septins partially depend on the exocyst for localization. **(A)** Co-localization of Spn1-mEGFP and Sec3-tdTomato at the division site in cells without (early) and with (late) septa. Sad1-mRFP1 marks the spindle pole body (SPB). **(B)** Line scans showing Spn1 and Sec3 intensities at the division site along the cell long axis in septated cells as in (A). **(C)** SoRa (Super resolution by Optical Re-Assignment) confocal microscopy of cells expressing both Spn1-mEGFP and Exo70-tdTomato showing their perfect colocalization in the middle focal plane. **(D)** Time course and **(E)** quantification (in minutes) of Sec3 and Spn1 localizations and appearance timing at the division site. SPB separation is defined as time 0. **(F)** Localization of Spn1 (<u>Max</u> intensity projection, <u>Middle</u> focal plane, and <u>End-on</u> view of the division site) in WT and *sec3-913* cells grown at 36°C for 4 h. Yellow boxes, cells without septa; Red boxes, cells with septa. **(G)** Localization of Spn1 and the contractile-ring marker Rng8 in *sec8-1* cells grown at 36°C for 4 h. **(H)** Spn1 intensities at the division site in WT and *sec3-913* cells grown at 36°C for 4 h. Cells were grouped into no septum, forming septum, and closed septum stages. **, P < 0.01; ***, P < 0.001. **(I)** FRAP analyses (photobleached at time 0) of Spn1 at the division site in WT and *sec3-913* cells grown at 36°C for 4 h. Mean ± SD. Bars, 5 μm.

Since the septin and exocyst colocalize at the division site and Sec3 arrives earlier, we tested whether septin localization depends on Sec3 and other exocyst subunits. In WT cells, Spn1 always formed ring structures at the rim of division plane during septation (Figure 1F). In exocyst mutants *exo70*Δ and the temperature sensitive *sec3-913* and *sec8-1*, Spn1 localization was comparable to WT at permissive temperature (Figure supplement S1A). At the restrictive temperature, although Spn1 localized as a ring at the division site before septation, a fraction of Spn1 abnormally spread onto the division plane following furrow ingression in *sec3-913* and *sec8-1* mutants (Figure 1F, red boxes; and Figure supplement S1B, middle focal plane). As *exo70*Δ cells have no severe defects (Wang et al., 2003), the exocyst complex may not be as compromised as in *sec3-913* and *sec8-1* mutants. Only minor mislocalization of Spn1 was observed in *exo70*Δ cells even at 36°C (Figure supplement S1B). This localization pattern in exocyst mutants suggested a possible correlation between septins and the furrow ingression. Indeed, some Spn1 followed the contractile ring marked with Rng8 (Wang et al., 2014) as it constricted, spread onto the new plasma membrane, and concentrated at the center of the division plane while maintaining its localization at the rim (Figure 1G). We also examined Spn1 levels at the division site in cells with no visible septum, forming septum, and closed septum. Spn1 levels were comparable or higher in exocyst mutants compared to WT at both 25°C and 36°C (Figures 1H and Figure supplement S1, C and D). FRAP analyses of Spn1 showed no difference in its dynamics in WT and *sec3-913* cells at 36°C (Figure 1I and Figure supplement S1E). Collectively, despite some Spn1 mislocalizes to the center of the division plane in exocyst mutants, majority of Spn1 still localizes to the rim. Thus, septins only partially depend on the exocyst for their localization.

Next, we examined the localization and levels of the exocyst complex (subunits Sec3, Exo70, and Sec8) in *spn1*Δ cells. In mitotic cells without a septum, the exocyst localized to the rim of the division plane in both WT and *spn1*Δ cells (Figures 2A and Figure supplement S1, F and G, yellow boxes). During septation, however, the exocyst spread across the division plane as a disk in *spn1*Δ cells while it remained at the rim in WT cells (Figures 2A and Figure supplement S1, F and G, red boxes). The levels of Sec3, Exo70, and Sec8 at the division site in *spn1*Δ cells were not significantly different from WT before septation (Figures 2B and Figure supplement S1H). During and after septum formation, the levels of all three exocyst components were significantly reduced at the division site (except Exo70 in cells with forming septum) and almost absent at the rim in *spn1*Δ cells (Figures 2, A and B; and Figure supplement S1, F-H). These results were confirmed in cells expressed Sec8-GFP and myosin light chain Rlc1 as a contractile ring marker (Figure 2, D and E). However, the dynamics of Sec3 at the division site was not affected in *spn1*Δ cells (Figures 2C and Figure supplement S1I). The exocyst was much more dynamic than septins at the division site (Figures 1H and 2C), which was confirmed by high temporal resolution imaging of Exo70 (Videos 2 and 3). Together, loss of septins results in exocyst mislocalization and its decreased levels, especially at the rim of the division plane during septum formation, suggesting that septins play an important role in regulating the exocyst localization at the division site.

**Figure 2.**
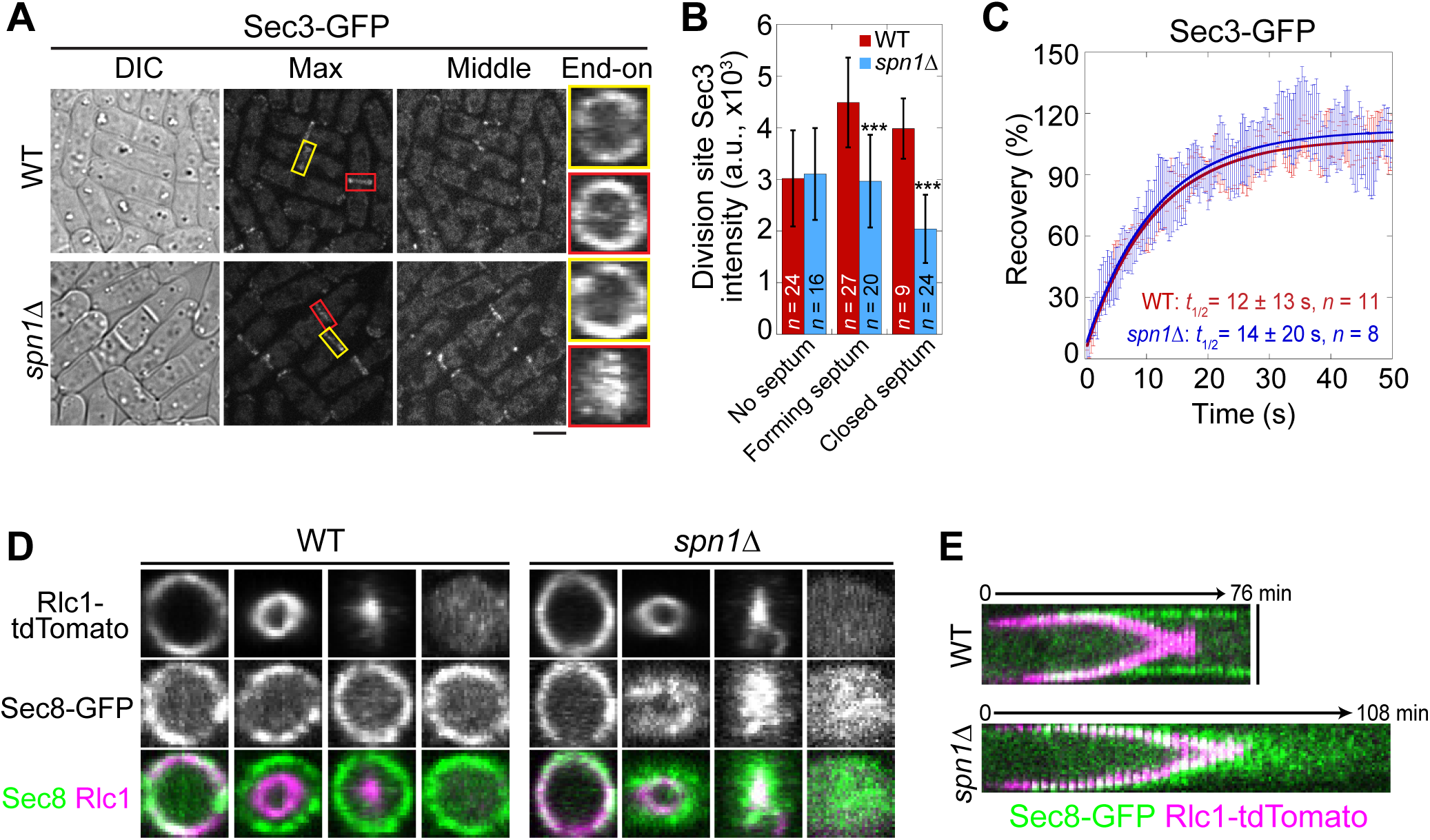
Septin rings recruit or anchor the exocyst complex to the rim of the division plane during late stage of cytokinesis. **(A)** Localization of Sec3 at the division site in WT and *spn1*Δ cells. Yellow boxes, cells without a septum; Red boxes, cells with a closed septum. **(B)** Sec3 intensity at the division site in WT and *spn1*Δ cells. ***, P < 0.001. **(C)** FRAP analyses of Sec3 at the division site in WT and *spn1*Δ cells. Mean ± SEM. **(D** and **E)** End-on views (D) and kymographs (E) of Sec8 and the contractile ring marker Rlc1 at the division site in WT and *spn1*Δ cells. Bars, 5 μm.

Collectively, our data suggest that septins and the exocyst complex colocalize and are interdependent for localization at the division site during and after the contractile ring constriction, with the septin rings being more important for the exocyst localization. Thus, we conclude that the initial recruitment of the exocyst to the division site does not depend on septins, but its rim localization and maintenance during late cytokinesis require the septin rings.

### Septins regulate the exocyst localization through physical interactions

Septins have been shown to play a role in the Rho GEF Gef3-Rho4 GTPase pathway to regulate the exocytosis of glucanases Eng1 and Agn1 for proper cell separation (Perez et al., 2015; Wang et al., 2015). Septins are essential for Gef3 localization to the division site (Muñoz et al., 2014; Wang et al., 2015). In *spn1*Δ cells, Gef3 localization on the plasma membrane is abolished, and Rho4 localization in cells with a closed septum was significantly reduced (Wang et al., 2015). Since Rho4 can interact with both the exocyst complex and septins (Perez et al., 2015), we tested whether the altered exocyst localization pattern that we observed in septin mutants was through Gef3 and Rho4. Although partially mislocalized to the center of the division plane, majority of Sec3 still localized as a ring at the rim of division plane in *rho4*Δ, *gef3*Δ, and *rho4*Δ *gef3*Δ cells (Figure supplement S2A). Moreover, Spn1 ring localization was not affected in *rho4*Δ *gef3*Δ cells (Figure supplement S2B). Thus, the different localization patterns of the exocyst in *spn1*Δ and *rho4*Δ *gef3*Δ cells suggest that septins can regulate exocyst localization independent of Gef3 and Rho4.

To test the hypothesis that septins regulate the localization of the exocyst directly, we examined the physical interactions between septins and the exocyst subunits. Sec3 and Exo70 are the most important subunits for the targeting of the octameric exocyst to the plasma membrane (Boyd et al., 2004; He et al., 2007; Bendezu et al., 2012; Luo et al., 2014; Yue et al., 2017; Liu et al., 2018; Synek et al., 2021), and Spn1 and Spn4 are essential for septin localization and functions (An et al., 2004). Therefore, we first tested the interactions between Spn1-Sec3, Spn1-Exo70, Spn4-Sec3, and Spn4-Exo70 using co-immunoprecipitation of cell extracts from fission yeast. Surprisingly, no physical interactions were detected among the four proteins.

Then we utilized AlphaFold2_advanced ColabFold algorithm (Jumper et al., 2021; Mirdita et al., 2022), whose highly accurate predictions of protein structures have revolutionized structural biology, to predict the physical interactions between all 32 combinations of the four septins and eight exocyst subunits. For the modeling, the complete sequences of each subunit of septins and exocyst complex were used except Sec8. Sec8 subunit was analyzed in two fragments with overlapping sequence due to the 1,400 amino acids input sequence limitation of AlphaFold2_advanced. The generated models with the highest confidence are shown (Figure 3 and Figure supplement S3). Predicted interacting interface residues that defined as amino acids of two possible binding partners with distance ≤4 Å were calculated from the rank 1 predicted model (Yin and Pierce, 2023). Then the contact residues were further narrowed down by excluding the residues having pLDDT score <50. Based on the above analyses, we predicted the following top six interactions between the septin and exocyst subunits: Spn2 and Sec15 (Figure 3), Sec15 and Spn1, Sec6 and Spn1, Spn2 and Sec5, Spn4 and Sec15, and Spn4 and Sec3 (Figure supplement S3).

**Figure 3.**
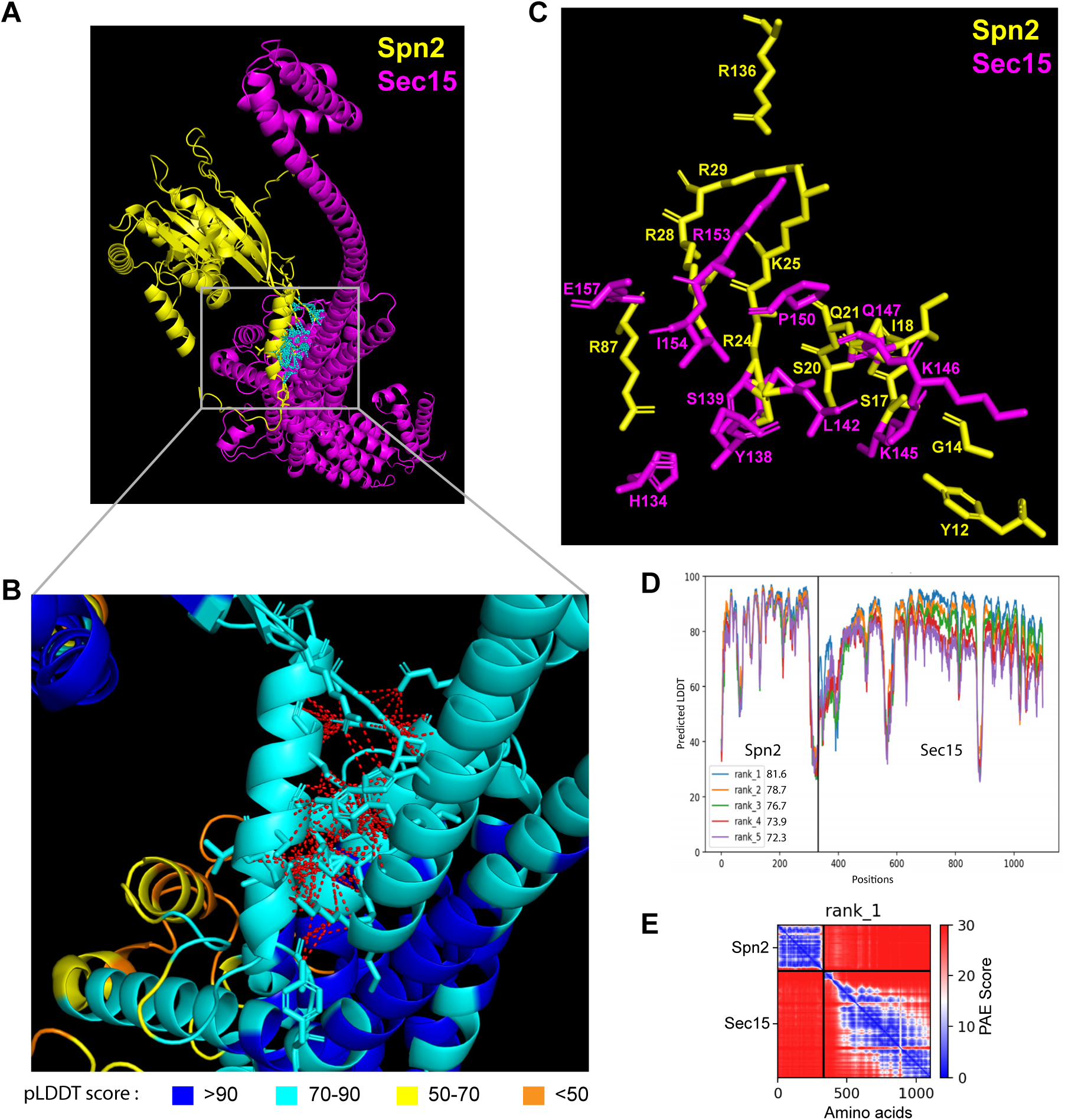
The 3D structural model of predicted interactions between Spn2 and Sec15 generated by AlphaFold. **(A-C)** AlphaFold2_advanced predicted interaction between Spn2 and Sec15 in rank 1 model with pLDDT score of 81.6. The pTM value = 0.51. Spn2 is colored in yellow and Sec15 in magenta in (A) and (C). **(B)** Inset of enlarged view of the predicted interactions, contacts between interface residues with distance <4 Å are colored in red (those in cyan in A). Residues are colored corresponding to their pLDDT scores as indicated in the legends below. (**C**) The key interacting residues and their side chains are shown. **(D)** Residue position scores of five predicted models for Spn2 and Sec15 interactions ranked according to pLDDT scores. **(E)** PAE (Predicted Alignment Error) plot for the top ranked model shown in (A-D), where colors represent confidence in the relative positioning of residues across the two proteins. Lower values (blue) represent high confidence while higher values (red) show low confidence in domain-domain interactions.

To test whether the interacting residues calculated from the pair-wise predictions are accessible in the whole exocyst or septin complexes, we employed AlphaFold3 and the cryo-EM structure of the whole *Saccharomyces cerevisiae* exocyst complex (with 4.4 Å resolution) to examine the interfaces of septins and the exocyst interactions, assuming that the *S. pombe* exocyst has the similar structure (Mei et al., 2018). For the septin complex, we first predicted its octameric structure using two copies of each fission yeast Spn1-4. We examined all the interacting residues on the septin and exocyst complex predicted from our AlphaFold2 modeling to determine whether these predicted interactions are structurally compatible (Figure supplement S4 and S5A; Videos 4 and 5). Our analyses revealed that 84% of exocyst and 96% of septin predicted interacting residues were sterically feasible without disrupting the architecture of exocyst or septin complex (the residues highlighted in yellow in Figure supplement S4 and S5; Videos 4 and 5), while others would likely require partial disassembly or flexible conformations. Because septins can also form hexameric complex (McMurray and Thorner, 2019; Mendonça et al., 2019; Soroor et al., 2021), we also predicted hexameric septin complexes without either Spn2 or Spn3 and found that 86% or 92% exocyst-interacting residues are available on the surface, respectively (Figure supplementary S5, B and C; Videos 6-7). We did not predict a septin hexameric complex without Spn1 or Spn4 as they are the core subunits responsible for septin localization and functions at the division site (An et al., 2004, Onishi et al., 2010; Wu et al., 2010). These predictions indicate that these septin-exocyst interactions are sterically plausible.

Next, we used reciprocal Co-IP assays of fission yeast extracts to confirm the predicted interactions between septin and exocyst subunits. Out of the six predicted interactions, we found five of them were positive in Co-IP. We found that Spn2 physically interacted with Sec5 and Sec15, Spn1 with Sec15 and Sec6, and Spn4 with Sec15 (Figures 4 and Figure supplement S6). Sec15 interacted with three septins Spn1, Spn2, and Spn4, which was stronger than other combinations. We also utilized yeast two-hybrid assays to confirm these five-pair of interactions (Figure 4, E and F). X-gal overlay assay (insets) and quantification of β-galactosidase using ONPG confirmed that Sec15 may directly interacted with Spn1, Spn2, and Spn4 (Figure 4E); and Sec6 interacted with Spn1 through its C-terminal fragment (Spn1[300-469]) that contains the coiled-coil motif (Figure 4F). The Spn2-Sec5 interaction could not be tested due to very high level of autoactivation of Sec5. Thus, we conclude that septins physically and likely interact directly with the exocyst in fission yeast via multivalent interactions.

**Figure 4.**
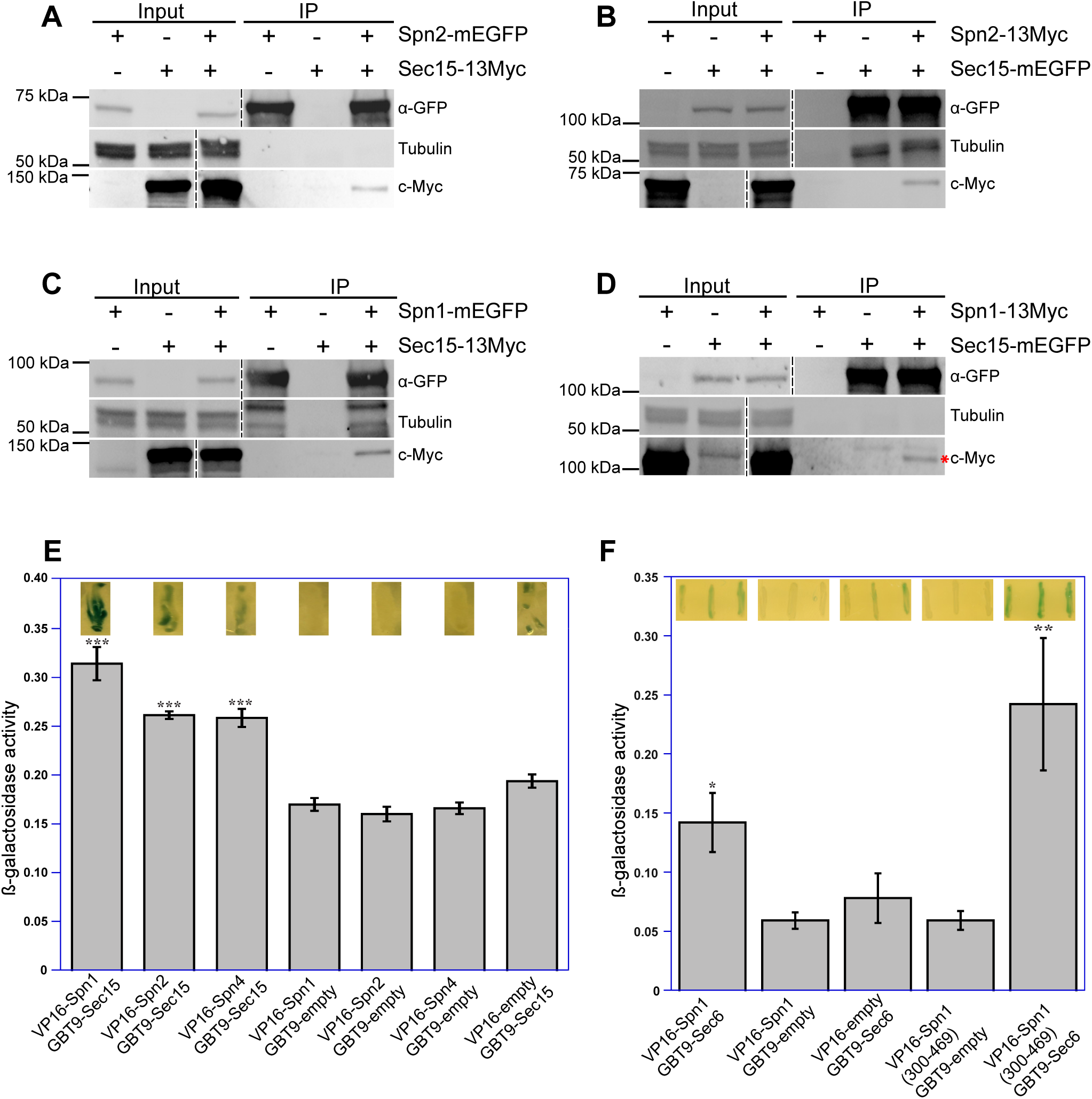
Septins and the exocyst interact physically. Reciprocal co-immunoprecipitation of Sec15 with Spn2 **(A** and **B)** and Spn1 **(C** and **D)**. Septin or exocyst subunits tagged with mEGFP or 13Myc were immunoprecipitated using antibodies against GFP from cell lysates, separated on SDS-PAGE, and incubated with appropriate antibodies. Tubulin was used as a loading control. Asterisk (*) in (D) marks Spn1-13Myc. The vertical dashed lines mark the positions of protein ladders that were excised out. (**E** and **F**) Septins and the exocyst subunits may interact directly revealed by the yeast two-hybrid assays. X-gal overlay results (insets on the top of the columns) and quantification of β-galactosidase activities using ONPG showing interactions between (E) Sec15 with Spn1, Spn2, and Spn4; and (F) Sec6 with Spn1 and its coil-coil motif Spn1(300-469). Data is shown in Mean ± SD, n = 3 (in E) or 4 (in F). ***p ≤ 0.0001, **p ≤ 0.001, *p ≤ 0.01 compared with their respective controls in one-way ANOVA with Tukey’s post hoc test.

### Septins are involved in concentrating Sec15 and Sec5 at the rim of the division plane especially during the late-stage of cytokinesis

We reasoned that septins localize the exocyst at the division site via their multivalent interactions with the exocyst subunits Sec15, Sec5, and Sec6. Weakened interactions between septins and the exocyst in the absence of a certain septin subunit could lead to mislocalization of the exocyst complex. Indeed, similar to the results presented in Figures 1, 2, and Figure supplement S1 with other exocyst subunits, the deletion of *spn1* or *spn4* led to mislocalization of Sec15 on the division plane in ∼75% of cells with a septum while Sec15 in ∼90% of WT cells localized as rings at the rim of the division plane in septating cells (Figure 5, A and B). Results from time-lapse microscopy of *spn1Δ* or *spn4Δ* cells were consistent with these findings. Sec15 was first recruited to division site as rings and then spread to whole division plane before signal disappearance, leading to some multiseptated cells (Videos 8-10).

**Figure 5.**
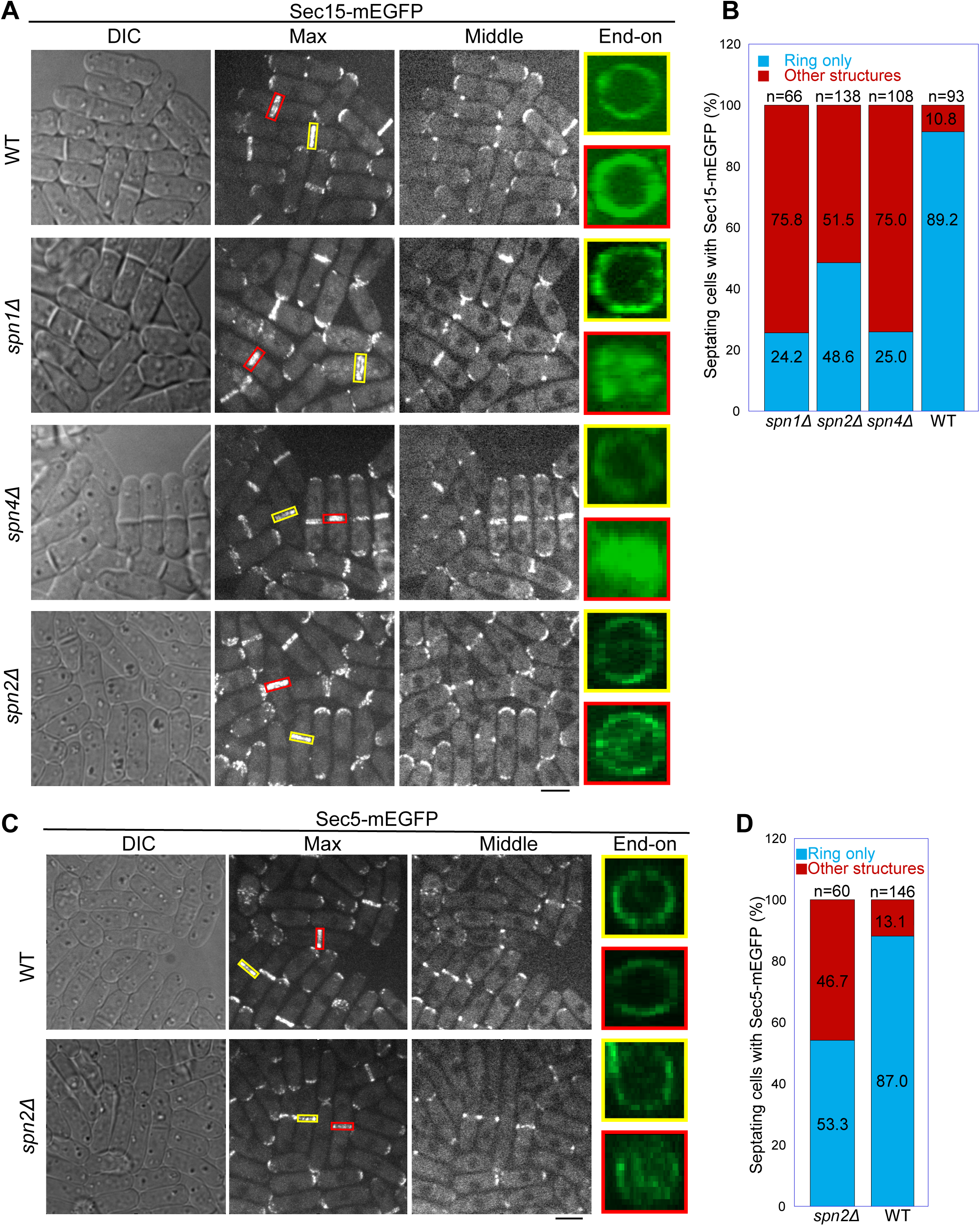
Localization patterns of both Sec15 and Sec5 at the division site depend on septins. Localization of **(A, B)** Sec15 and **(C, D)** Sec5 at the division site in WT and septin mutant cells. Yellow boxes, cells without a septum; Red boxes, cells with a closed septum in (A, C). (B, D) Quantification of cells with intact and mislocalized Sec15 (B) and Sec5 (D) signals in WT and septin mutant cells with obvious septa. Scale bars, 5 µm.

Although septin filaments have four subunits in vegetative cells (An et al., 2004), Spn2 is less important than Spn1 and Spn4 for septin functions, and *spn2Δ* has a much weaker phenotype in septation than *spn1Δ* or *spn4Δ* (An et al., 2004; Wu et al., 2010; Zheng et al., 2018). Consistently, in *spn2Δ* cells, Sec15 and Sec5 localized normally at the division site before septation (Figure 5, A and C). Both Sec15 and Sec5 spread more or less to the whole division plane in ∼50% of *spn2Δ* cells with obvious septa in the DIC channel (Figure 5, A-D). Unlike in *spn1Δ* or *spn4Δ* cells, a fraction of Sec15 and Sec5 can still localize to the rim in *spn2Δ* cells (Figure 5, A and C). Collectively, these data support that Spn1, Spn2, and Spn4 are important for restricting the exocyst to the rim of division plane during cytokinesis through physical interactions.

### Septin mutants affect the sites of secretory vesicle tethering and cargo delivery at the division plane

Septin and exocyst mutations showed no or very mild synthetic genetic interactions (Tables 1 and 2), suggesting that septins and the exocyst complex function in the same pathway to regulate cytokinesis and septation. Surprisingly, they have different genetic interactions with the transport particle protein-II (TRAPP-II) mutants (Tables 1 and 2). The exocyst mutant *sec8-1* is synthetic lethal with *trs120-M1* and has severe synthetic cytokinesis defects with *trs120-ts1* due to the overlapping function of the exocyst and TRAPP-II in exocytosis during fission yeast cytokinesis (Wang et al., 2016). However, *spn1*Δ *trs120-M1* and *spn1*Δ *trs120-ts1* double mutants were viable with no obvious synthetic interactions (Tables 1 and 2). Thus, septins and the exocyst also work in different genetic pathways for certain functions in fission yeast.

**Table 1.**
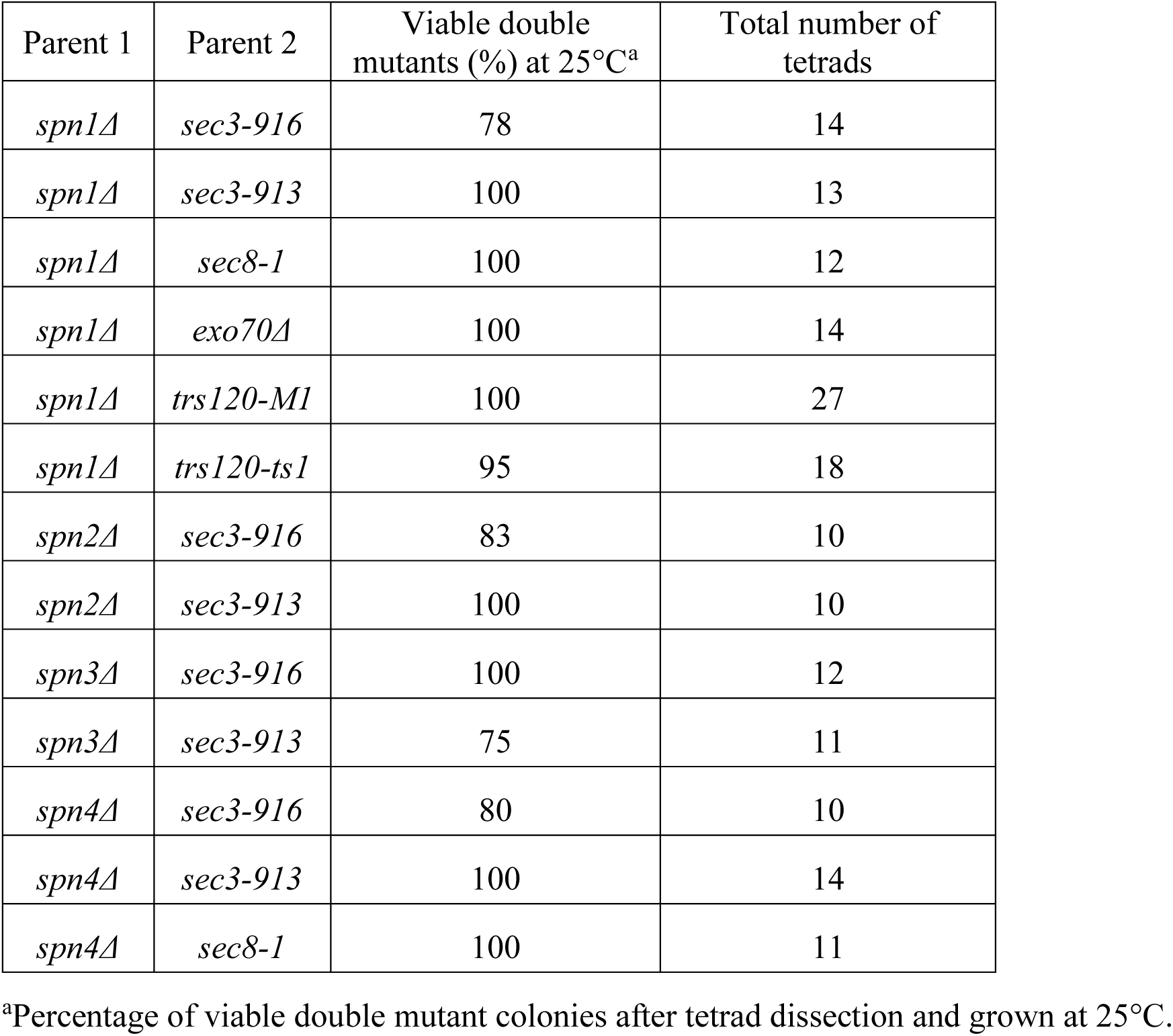
Viability of double mutants of the septin and exocyst from tetrad dissection at 25°C.

**Table 2:** Genetic interactions between septin and exocyst mutations at various temperatures.^a^.

| Mutations | 25°C | 30°C | 32°C | 36°C |
| --- | --- | --- | --- | --- |
| *sec3-916* | +++<sup>b</sup> | ++<sup>c</sup> | +<sup>d</sup> | -<sup>e</sup> |
| *sec3-913* | +++ | +++ | ++ | - |
| *sec8-1* | +++ | ++ | + | - |
| *spn1Δ* | ++ | ++ | ++ | ++ |
| *spn1Δ sec3-916* | ++ | + | - | - |
| *spn1Δ sec3-913* | ++ | ++ | ++ | - |
| *spn1Δ sec8-1* | ++ | ++ | + | - |
| *exo70Δ* | +++ | +++ | ++ | - |
| *spn1Δ exo70Δ* | ++ | ++ | ++ | - |
| *trs120-M1* | ++ | - | - | - |
| *spn1Δ trs120-M1* | ++ | - | - | - |
| *trs120-ts1* | +++ | +++ | + | - |
| *spn1Δ trs120-ts1* | ++ | ++ | + | - |
| *spn2Δ* | +++ | +++ | +++ | +++ |
| *spn2Δ sec3-916* | ++ | + | + | - |
| *spn2Δ sec3-913* | +++ | ++ | + | - |
| *spn3Δ* | +++ | +++ | +++ | +++ |
| *spn3Δ sec3-916* | ++ | ++ | + | - |
| *spn3Δ sec3-913* | +++ | ++ | ++ | - |
| *spn4Δ* | ++ | ++ | ++ | ++ |
| *spn4Δ sec3-916* | ++ | + | - | - |
| *spn4Δ sec3-913* | ++ | ++ | ++ | - |
| *spn4Δ sec8-1* | ++ | ++ | ++ | - |

<sup>a</sup>Cells were freshly grown on YE5S and YE5S plus Phloxin B (PB, which accumulates in dead cells) plates before checking the growth and morphology under DIC at different temperatures. The defects in cytokinesis and cell integrity compared with the parent strains was classified as follows:
<sup>b</sup>+++ , comparable to wt.
<sup>c</sup>++ , some cell lysis or cytokinesis defects.
<sup>d</sup>+ , severe cytokinesis defects with reduced growth.
<sup>e</sup>- , inviable.

The exocyst complex is the major tether of secretory vesicles at the plasma membrane (TerBush and Novick, 1995; TerBush et al., 1996; Wang et al., 2002; Luo et al., 2014). So we tested whether exocyst mislocalization in septin mutants compromises the targeting of secretory vesicles and their cargos. We first performed electron microscopy to examine if secretory vesicles are accumulated at the division site in *spn1*Δ cells (Figure 6A). During septum formation, 7- and 2-fold more secretory vesicles accumulated at the division site in *sec8-1* and *spn1*Δ cells, respectively, compared to WT (Figure 6, A and B). However, in cells with a closed septum, the number of secretory vesicles adjacent to the division site were not significantly different between WT and *spn1*Δ cells (Figure 6B). Consistently, secretory vesicle markers Rab11 GTPase Ypt3 and v-SNARE Syb1 accumulated more in the center of the division plane but diminished from the rim in *spn1*Δ cells (Figure 6, C and D). The accumulation of the secretory vesicles at the division plane and their mistargeting are consistent with exocyst mislocalization in *spn1*Δ cells.

**Figure 6.**
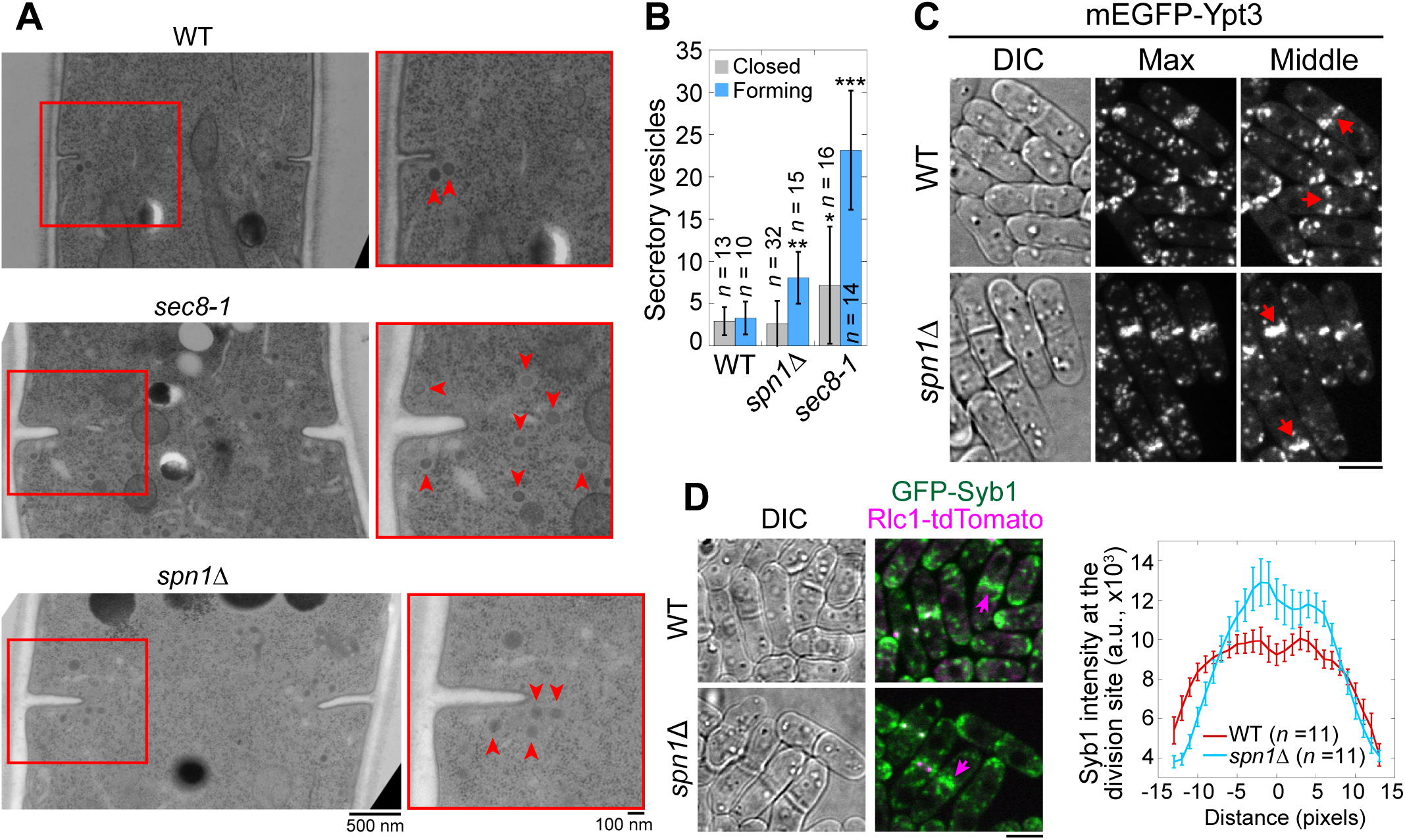
Septins are important for proper localization and distribution of secretory vesicles. **(A** and **B)** EM thin-section images (A) and quantifications of secretory vesicles (B) in WT, *sec8-1*, and *spn1*Δ cells with forming or closed septa. Cells were grown at 36°C for 4 h. Red boxes indicate the enlarged regions on the right. Arrowheads mark secretory vesicles. *, P < 0.05; **, P < 0.001; ***, P < 0.0001 compared to WT. *n* = numbers of thin sections. **(C** and **D)** Localizations of the Rab11 GTPase Ypt3 (C) and the v-SNARE Syb1 and Rlc1 (D) in WT and *spn1*Δ cells. Arrows mark examples of cells with closed septa. Syb1 intensities at the division site (D, right) from line scans at the middle focal plane of cells with closed septa (at the end of ring constriction indicated by an Rlc1 dot at the center of the division plane). Bars, 500 nm (A, left), 100 nm (A, right), and 5 μm (C and D).

We next examined the distribution of two secretory vesicle cargos, β-glucan synthase Bgs1/Cps1 and β-glucanase Eng1, which were delivered to the division site by the secretory vesicles during cytokinesis (Liu et al., 1999; Baladrón et al., 2002; Cortes et al., 2002; Martin-Cuadrado et al., 2003). More Bgs1 localized in the center of the division plane in *spn1*Δ cells compared to WT (Figure 7A). *spn1*Δ and *sec8-1* cells also had thicker septa compared to WT cells (Figure 7B). Another cargo of secretory vesicles, Eng1, spread across the division plane as a disk with localization clearly missing at the rim in *spn1*Δ cells (Figure 7C). Lack of glucanase Eng1 at the rim could contribute to the delayed cell separation in *spn1*Δ cells since the junctions between septum and the cell wall cannot be efficiently digested, consistent with earlier studies (Baladrón et al., 2002; Martin-Cuadrado et al., 2003). Our studies on Bgs1 and Eng1 indicate an increase of vesicle tethering in the center and a loss at the rim of the division plane without septins.

**Figure 7.**
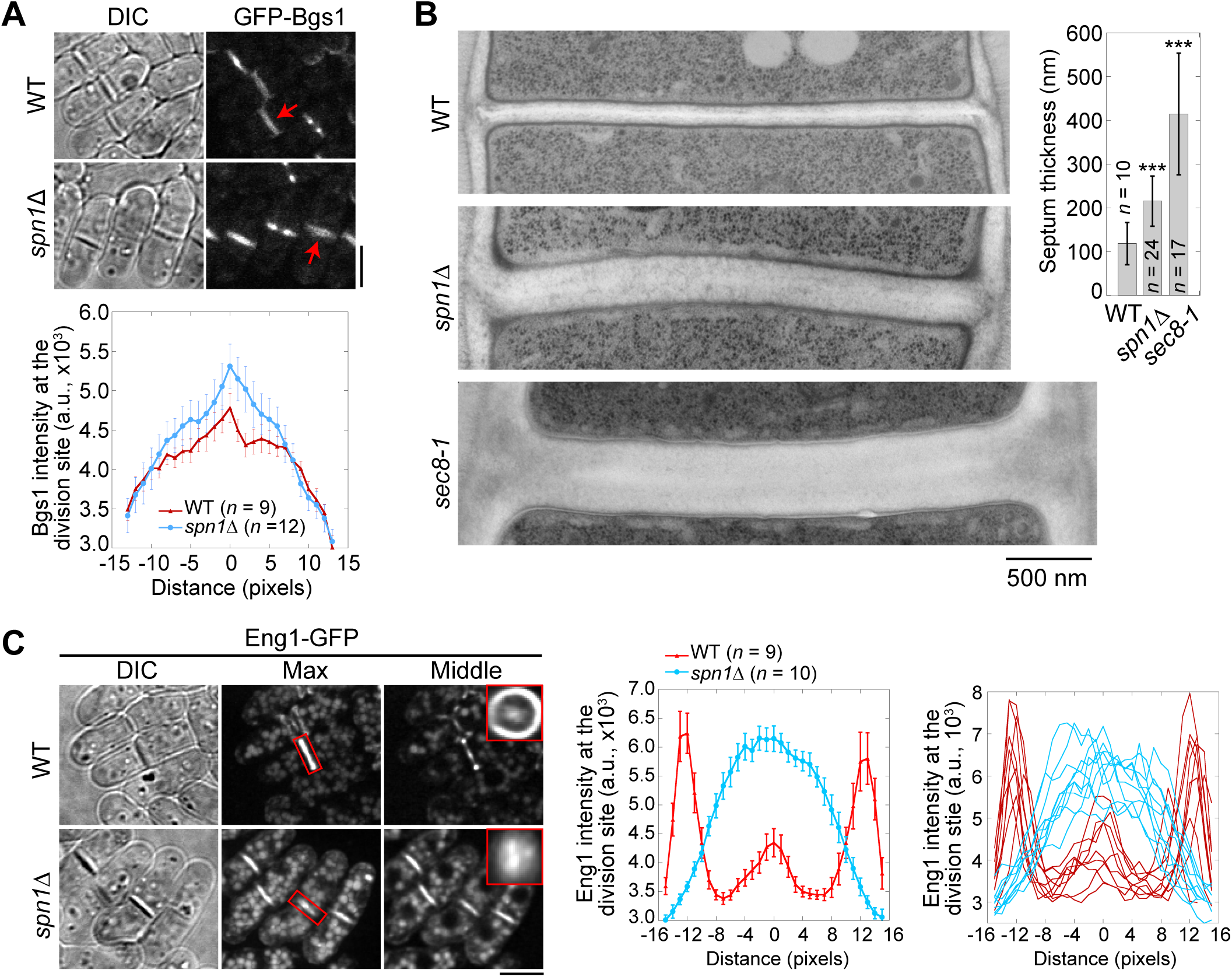
Septins are important for localization and distribution of secretory cargos Bgs1 and Eng1. **(A)** Localization (top) and intensity (bottom) of the glucan synthase Bgs1 in WT and *spn1*Δ cells. Arrows mark examples of cells with a closed septum. Bgs1 intensities from line scans across the division site at the middle focal plane were compared in cells with closed septa. **(B)** EM thin-section images (left) and septum thickness (right) of WT, *spn1*Δ, and *sec8-1* cells with closed septa. Cells were grown at 36°C for 4 h. ***, P < 0.0001 compared to WT. **(C)** Localization (left) and intensity (middle and right) of Eng1-GFP in WT and *spn1*Δ cells. The end-on views of Eng1 at the division site in cells with closed septa are shown as insets. Eng1 intensities (Middle, mean intensities; and Right, individual cells) are from line scans at the middle focal plane. Bars, 5 μm (A and C) and 500 nm (B).

Collectively, our data indicate that septins play important roles in maintaining the proper localization and targeting of the exocyst on the plasma membrane. This regulation on the exocyst occurs specifically during late stages of cytokinesis in fission yeast. Loss of septins results in spreading of the exocyst across the division plane and tethering of secretory vesicles at wrong destination, which leads to the accumulation of secretory vesicles and mistargeting of downstream cargos.

## Discussion

In this study, we reveal that septins and the exocyst complex physically interact to regulate exocytosis and ensure proper targeting of vesicle cargos to the plasma membrane during cytokinesis.

### Septins are important for proper membrane targeting of the exocyst complex to ensure successful cytokinesis

Septins are essential for cytokinesis and other cellular processes in budding yeast and many other organisms (Neufeld and Rubin, 1994; Longtine et al., 1996; Kinoshita et al., 1997; Gladfelter et al., 2001; Russell and Hall, 2005; Oh and Bi, 2011). However, the nature of their functions is only partially understood. It has been a mystery why the phenotypes of septin mutants are quite mild in fission yeast ever since their discoveries in the early 1990s, yet their sequences and structures are evolutionarily conserved across species (Longtine et al., 1996; An et al., 2004; Zheng et al., 2018; Zheng et al., 2024). In this study, we investigated the spatial regulation of the exocyst complex and roles of septins during cytokinesis in the fission yeast model system. Without septin rings, the exocyst complex, which specifies for the sites for vesicle fusion on the plasma membrane, cannot maintain its localization at the rim of the division plane. Instead, the exocyst complex follows actomyosin contractile ring constriction and spreads across the whole division plane. Although loss of septins does not affect the dynamics of the exocyst, the targeting sites of secretory vesicles and their cargos are altered, which may contribute to a thicker septum and a delayed cell separation. The modest accumulation of vesicles and vesicle cargos at the division site is one of the reasons for the increased thickness of the division septum in septin mutants. It is more likely that the misplaced exocyst can still tether vesicles along the division plane without septins. Due to the lack of the glucanase Eng1 at the rim of the division plane in septin mutants, daughter-cell separation is delayed and then cells continue to thicken the septum. The relatively modest vesicle accumulation in septin mutants suggests that septins are not absolutely required for vesicle tethering or fusion per se at the division site. Instead, septins primarily function to spatially organize the targeting sites of exocyst-directed vesicles by stabilizing the localization of the exocyst at the rim of the cleavage furrow. In septin mutants, mislocalization of the exocyst reduces the spatial precision of membrane insertion but still permits vesicle tethering and fusion, albeit in a less controlled manner. Thus, septins likely play a modulatory rather than essential role in exocytic vesicle delivery during cytokinesis. This interpretation aligns with our localization and genetic interaction data, which indicates that septins act as scaffolds to optimize secretion geometry, rather than as core components of the fusion machinery. Thus, fission yeast septins function in exocytosis through maintaining proper docking sites of the exocyst complex and secretory vesicles at the division site.

Both the exocyst and TRAPP-II complex tether vesicles at the cleavage furrow during cytokinesis (Wang et al., 2016). The genetic interactions between mutations in the exocyst and septins when combined with TRAPP-II mutants may reflect fundamentally different consequences for compromising the exocyst function (Tables 1 and 2). In septin mutants, the exocyst complex still localizes to the division site but is mispositioned from the rim to the center of the division plane. This mislocalization allows partial retention of exocyst function, leading to very mild synthetic or additive defects when combined with compromised TRAPP-II trafficking and tethering. In contrast, in exocyst subunit mutants, the exocyst becomes partial or non-functional, resulting in a more severe loss of exocyst activity. These differing consequences could explain the qualitative differences in genetic interactions observed with TRAPP-II mutants (Tables 1 and 2). Thus, septins and the exocyst also work in different genetic pathways for certain functions in fission yeast cytokinesis.

Fission yeast septins regulate the exocyst in specific temporal and spatial manners. They only regulate the localization of the exocyst during late stages of cytokinesis and are not responsible for its targeting to the cell tips during interphase or initial recruitment to the division site during early cytokinesis before septin appearance (Figures 1D, 2A; Figure supplement S1, F and G; and Videos 8-10). Disruption of the contractile ring affects the localization of the exocyst to the division site (Wang et al., 2002; Dobbelaere and Barral, 2004). This suggests that the exocyst likely depends on the contractile ring components for initial recruitment to the division site. However, this is not a universal mechanism. The subcellular localization of the exocyst complex in rat brain cells is affected by microtubule, but not actin-disrupting drugs (Vega and Hsu, 2003). Thus, how the exocyst is initially recruited to the division site remains to be studied. Since fission yeast exocyst clearly depends on septins for proper localization during late stages of cytokinesis, its localization dependence must migrate to septin rings from the contractile ring at some point before the onset of the contractile ring constriction. So it will be of great interest to examine how this transition occurs. Although septins may act as either scaffolds or diffusion barriers for Sec3 in budding yeast, Sec3 localizes between the split septin rings during cytokinesis (Dobbelaere and Barral, 2004). However, in mammalian neurons, the exocyst subunits Sec6 and Sec8 colocalize with the septin SEPT7/CDC10 (Hsu et al., 1998). Thus, the colocalized septins and the exocyst in fission yeast may provide more insights in mammalian cells for understanding the molecular mechanisms of their interactions.

Examples of localization dependence between septins and the exocyst have been reported in other systems. The most prominent cases come from fungal pathogens (Eisermann et al., 2023). *Magnaporthe oryzae* infects plants through a specialized infection cell called appressorium, which breaches through the cuticle of the leaf to allow entry into plant tissues (Dagdas et al., 2012; Gupta et al., 2015; Zhang et al., 2021). The exocyst assembles in appressorium at the point of plant infection in a septin-dependent manner. Septin deletion causes mislocalization of the key component for the exocyst assembly, Sec6, at the appressorium pore (Gupta et al., 2015). Similarly, the root-infecting phytopathogenic fungus *Verticillium dahliae* also assembles the exocyst at the penetration peg of the hyphopodium in a septin-dependent manner (Zhou et al., 2017). The absence of septin VdSep5 impairs the delivery of secretory proteins to the penetration interface (Zhou et al., 2017). Another example is *Candida albicans* septins, which localize at the hyphal tips where tip growth occurs with active exocytosis in this human opportunistic pathogen (Li et al., 2007). Deletion of septin *CDC10* or *CDC11* causes mislocalization of the exocyst marked by Sec3 (Li et al., 2007). Thus, one of the conserved roles of septins is to regulate the proper membrane targeting of the exocyst complex to the plasma membrane and to ensure spatiotemporal fidelity of vesicle tethering and fusion. Our current study will provide insights into how septins and the exocyst help fungal pathogens infect their hosts. However, how they physically interact with each other had not been systematically investigated.

### The exocyst complex docks on septins on the plasma membrane through multivalent physical interactions

Despite the relationships between septins and the exocyst mentioned above, whether and how they physically interact with each other were obscure. In budding yeast, the exocyst subunits have been shown to interact physically with a number of proteins, including Sec15 with Rab GTPase Sec4 and type V myosin Myo2; and Sec6 with v-SNARE protein Snc2, t-SNARE protein Sec9, and Sec1/Munc18 family protein Sec1 (Guo et al., 1999; Sivaram et al., 2005; Jin et al., 2011; Shen et al., 2013; Lepore et al., 2016). The septin dynamics is essential for exocytosis (Tokhtaeva et al., 2015). But septins and the exocyst do not colocalize in budding yeast (Dobbelaere and Barral, 2004; Okada et al., 2013). Active Cdc42 recruits septins to the polarization site. The septin ring that is formed by polarized exocytosis corrals exocyst-dependent exocytosis and active Cdc42 inside the ring (Dobbelaere and Barral, 2004; Okada et al., 2013). However, there is no evidence that the exocyst and septins physically and directly interact in budding yeast. Consistently, recent mapped *S. cerevisiae* protein interactome found no interactions between septins and exocyst in the pull down experiments (Michaelis et al., 2023).

By contrast, several interactions between septins and the exocyst have been identified by Co-IPs to support the role of septins in the regulation of the exocyst localization in other cell types (Hsu et al., 1998; Beites et al., 1999; Vega and Hsu, 2003; Li et al., 2007; Gupta et al., 2015). In rat brain lysates, the septins SEPT2, 4, 6, and 7 have been shown to associate with the exocyst complex containing Sec6/8 (Hsu et al., 1998). Sec6 is found to be colocalized with SEPT7 on synapse assembly sites in isolated neurons where active membrane remodeling is required (Hsu et al., 1998). From rat brain cells, the exocyst subunits Sec8 and Exo70 along with tubulin co-immunoprecipitated with the septin Nedd5 (Vega and Hsu, 2003). During hyphal development in *C. albicans*, association of Sec3 and Sec5 with the septin Cdc3 was detected by co-IP (Li et al., 2007). In *M. oryzae,* mislocalization of Sec6 was reported with deletion of the septin Sep3. This was supported by pull down and mass spectrometry data where Sep4 and Sep5 were pulled down by Exo84 while Sep3 by Sec6 (Gupta et al., 2015). Here we have presented comprehensive studies on explaining importance of septins in regulating exocytosis by likely direct physical interactions with the exocyst in fission yeast.

In our study, we systematically investigated all the potential pairwise interactions between septin and exocyst subunits using AlphFold2 predictions. We experimentally confirmed five out of the six predicted interactions by co-IPs: Spn1-Sec15, Spn1-Sec6, Spn2-Sec15, Spn2-Sec5, and Spn4-Sec15 and validated four of them by yeast two-hybrid assays (except Spn2-Sec5 due to high levels of Sec15 autoactivation). The observed associations are consistent with direct interactions predicted by AlphaFold2, but cannot alone establish their direct bindings. These multivalent interactions ensure that the exocyst dynamically tethers secretory vesicles on the plasma membrane with high temporal and spatial fidelity even individual interaction may not be very strong. The subunits Sec15, Sec6, and Sec5 in the exocyst complex are known to be available for interacting with many proteins as mentioned above in budding yeast and in other systems for different cellular functions (Sjölinder et al., 2002; Fukai et al., 2003; Zhang et al., 2004; Feng et al., 2012; Du et al., 2015; Guo et al., 2016; Yang et al., 2022). We also predicted the whole *S. pombe* exocyst structure using AlphaFold3, which is most likely based on the published structural models of the full exocyst complex from budding yeast (e.g., PDB: 5YFP; Lepore et al., 2018; Mei et al., 2018). We also predicted the structures of septin octameric and hexameric complexes. We found that majority of the predicted exocyst-septin interacting residues are located on the accessible surfaces of the assembled whole complexes (Figure supplement S4 and S5; and Videos 4-7). These predictions indicate that these septin-exocyst interactions are sterically plausible. The interactions between septins and the exocyst that we identified in fission yeast will provide important insights into the mechanisms of exocyst regulations by septins. During evolution, fission yeast may have lost many but some of the most conserved aspects of septin functions including septin-exocyst interactions.

It is known that the octameric exocyst complex consists of two subcomplexes (Heider et al., 2016; Ahmed et al., 2018; Lepore et al., 2018; Mei and Guo, 2018; Mei et al., 2018; Ganesan et al., 2020). Subcomplex 1 consists of Sec3, Sec5, Sec6, and Sec8 while subcomplex 2 consists of Sec10, Sec15, Exo70, and Exo84. In our study, we found that septins can interact with both of the exocyst subcomplexes with multivalent interactions by Alphafold predictions, reciprocal Co-IPs, and yeast two-hybrid assays. Some of the identified interactions may only be strong enough between specific subunits at exposed interfaces under the Co-IP conditions, rather than through the whole complex as predicted by AlphaFold. Additionally, the detergent and salt conditions used in our co-IPs may disrupt labile complex interfaces or partially dissociate multimeric assemblies. Future studies are needed to refine the residues involved in the interactions because the predicted interacting residues from AlphaFold are too numerous. However, it is encouraging that most of the predicted interacting residues are clustered in several surface patches. Experimental validation through targeted mutagenesis is an important next step. In addition, tests are needed to figure out if posttranslational modifications are necessary for the interactions between septins and the exocyst. Because the colocalization of septins and the exocyst required for their proper function occurs at a specific stage during cytokinesis rather than a general regulation throughout the cell cycle, septin filament formation and posttranslational modifications of the involved proteins are most likely required, which make it challenging to tease out the interactions in vitro (Dobbelaere et al., 2003; Hernández-Rodríguez and Momany, 2012; Ren and Guo, 2012; Tay et al., 2019; Sharma and Menon, 2023; Werner and Yadav, 2023). Moreover, we cannot rule out that Rho1/RhoA GTPase and PI(4,5)P2 are involved in septin-exocyst interactions as both have been reported to interact with septins and/or the exocyst in other cell types (Guo et al., 2001; He et al., 2007; Bertin et al., 2010; Bendezu and Martin, 2011; Perez et al., 2015; Carim and Hickson, 2023; Safavian et al., 2023).

In summary, we found that septins are important for exocyst targeting to the division site during late stages of cytokinesis through multivalent interactions between their subunits. The proper exocyst localization at the rim of the division plane is critical for timely and successful cytokinesis. Our results will provide insights into future studies of the interactions and functions of both septins and the exocyst in other cell types. Dysregulation of septins or the exocyst leads to severe disorders including neurological diseases and cancers (Russell and Hall, 2005; Martin-Urdiroz et al., 2016; Halim et al., 2023; Werner and Yadav, 2023). Thus, it is important to identify the functional and physical links between septins and the exocyst complex in human cells.

## Materials and methods

### Strains and molecular methods

Fission yeast strains used in this study are listed in Supplemental Table S1. Strains were constructed using PCR-based gene targeting and standard genetic methods (Moreno et al., 1991; Bähler et al., 1998). Tagged genes were expressed under endogenous promoters and integrated at their native chromosomal loci except where noted. The glucan synthase gene *bgs1* is integrated at the *leu1* loci under endogenous promoter, with the endogenous copy deleted (Cortes et al., 2002). The functionalities of the newly tagged proteins (Spn1, Spn2, Spn4, Sec3, Sec5, Sec6, Sec8, Sec15, and Exo70) were tested by growing the strains at 25°C and 36°C on YE5S media or crossing to mutants. The growth and morphology of the tagged strains were comparable to WT.

### Microscopy

Cells were normally grown at the exponential phase in YE5S liquid medium at 25°C for 40-48 h before microscopy or temperature shift. Confocal microscopy was performed as previously described (Wang et al., 2014; Davidson et al., 2015; Davidson et al., 2016; Zhu et al., 2018). Briefly, cells were collected from liquid culture by centrifuging at 3,000 rpm for 30 s at room temperature and washed with EMM5S twice to reduce autofluorescence. A final concentration of 5 µM *n*-propyl-gallate (*n*-PG) from a 10x stock (in EMM5S) was added in the second wash to protect cells from free radicals during imaging. Live cells were imaged on a thin layer of EMM5S with 20% gelatin and 5 µM *n*-PG at ∼23°C. To image cells at 36°C, concentrated cells were spotted into coverglass-bottom dish and covered with EMM5S agar (Davidson et al., 2016). We imaged cells using several microscopy systems with 100x/1.4 or 100x/1.45 numerical aperture (NA) Plan-Apo objective lenses (Nikon, Melville, NY). Most fluorescence images were taken using a PerkinElmer spinning disk confocal system (UltraVIEW Vox CSUX1 system; PerkinElmer, Waltham, MA) with 440-, 488-, 515-, and 561-nm solid-state lasers and back thinned electron-multiplying charge-coupled device (EMCCD) cameras (C9100-13 or C9100-23B; Hamamatsu Photonics, Bridgewater, NJ) on a Nikon Ti-E inverted microscope. For better spatial resolution, Figure 1A was imaged using another spinning disk confocal system (UltraVIEW ERS; PerkinElmer) with 568-nm solid-state laser and 488-nm argon ion lasers and a cooled charge-coupled device camera without binning (ORCA-AG; Hamamatsu Photonics) on a Nikon Eclipse TE2000-U microscope. For the SoRa imaging shown in Figure 1C, the images were captured with a Nikon CSU-W1 SoRa spinning disk confocal system equipped with 488 and 561-nm solid-state lasers and an ORCA-Quest qCMOS camera (C15550, Hamamatsu Photonics, Bridgewater, NJ) on a Nikon Eclipse Ti-2E microscope with 2x2 binning (Ye et al., 2025). We used TIRF microscopy controlled by NIS Elements software to examine the dynamic localization of the exocyst subunit Exo70 and the septin Spn1 at the division site for some movies (Videos 1-3). A Nikon Eclipse Ti-E microscope equipped with a TIRF illuminator, Plan Apo 100x/1.45NA oil objective, and an Andor iXon Ultra 897 EMCCD was used.

### Image analysis

We analyzed images using ImageJ/Fiji (National Institutes of Health, Bethesda, MD) and Volocity (PerkinElmer). Fluorescence images are maximum-intensity projections from z-sections spaced at 0.5 μm except where noted. Images of 3D projections (end-on views) and deconvolution (Figure 7C, Eng1) were generated from images with z-sections spaced at 0.05 μm. For quantification of fluorescence intensity at the division site, we summed the intensity from all z-sections using sum projection. A rectangular ROI1 was drawn to include majority of division site signal for intensity measurement. Then the intensity in a second ROI2 approximately twice the area of ROI1 (including ROI1) was measured and used to subtract cytoplasmic background as described previously (Coffman et al., 2011; Davidson et al., 2015; Davidson et al., 2016).

For comparing the colocalization at the rim of the division plane (Figure 1B), a line along the cell long-axis was drawn across the division plane at the same position for both Spn1 and Sec3 channels using maximum intensity projection images. Then the width of the line was adjusted to cover all signals at the division site, generating an ROI of 1.5 μm x 3.5 μm (x-y) (see Figure 1B). The mean intensity of all pixels in y-axis was measured along x-axis and plotted.

Line scans (Figures 6D, 7A, and 7C) across the division plane were made in the middle focal plane of the fluorescence images. A line along the cell short-axis was drawn across the division plane of the cells with a closed septum (at or after the end of contractile-ring constriction) to cover the whole cell diameter. To quantify their fluorescence intensity at the division site using line scans, the line width used was 3 pixels to reduce signal variations caused by measurements on a single focal plane. Mean intensity (average of 3 pixels) was measured across cell diameter. For Syb1 (Figure 6D), cells at the end of ring constriction (indicated by an Rlc1 dot at the center of the division plane) were measured; and line scans were aligned by referencing the peak intensity of Rlc1 signal. All data were aligned by the center and plotted. For Bgs1 (Figure 7A), we quantified the cells that Rlc1 signal had disappeared from the division site. The line was drawn in the Bgs1 channel in the middle focal plane. The center of line scan was defined as the pixel with the brightest Bgs1 value. All data were aligned by the center and plotted. For Eng1 (Figure 7C), cells with closed septa were measured; and line scans for WT cells were aligned by the middle of the two peaks; and the ones for *spn1*Δ cells were aligned by referencing the middle of septa in DIC images.

### FRAP analysis

FRAP was performed using the photokinesis unit on the UltraVIEW Vox confocal system at either ∼23°C or 36°C (Coffman et al., 2009; Laporte et al., 2011; Zhu et al., 2013). Half of the division site signals at the middle focal plane were photobleached to <50% of the original fluorescence intensity. Five pre-bleach images and 150 post-bleach images for *spn1*Δ cells, or 70 post-bleach images for *sec3-913* cells, were collected at every 0.33 s or 10 s, respectively. For image analysis, the background and photobleaching during image acquisition were corrected using empty space and unbleached cells within the same image. The pre-bleach intensity was normalized to 100%, and the first post-bleach intensity was normalized to 0% (Laporte et al., 2011; Zhu et al., 2018). Intensities of three consecutive post-bleach time points were rolling averaged to reduce noise (Vavylonis et al., 2008). Data were plotted and fitted using the exponential decay equation *y* = *m*_1_ + *m*_2_ exp(-*m*_3_*x*), where *m*_3_ is the off-rate. The half-time for recovery was calculated by *t*_1/2_ = ln2/*m*_3_.

### Predictions of septin-exocyst interactions using AlphaFold analyses

The development of computer algorithms to predict three-dimensional protein structures from amino acid sequence involves two complementary ways that concentrate on either the physical interactions or the evolutionary history (Jumper et al., 2021). AlphaFold utilizes cutting-edge neural network topologies and training techniques to predict the 3D coordinates of a primary amino acid sequence (Jumper et al., 2021). We made the AlphaFold models of interactions between different septin and exocyst subunits using Google Colab Platform and AlphaFold2_advanced option that does not need templates at: https://colab.research.google.com/github/sokrypton/ColabFold/blob/main/beta/AlphaFold2_advanced.ipynb#scrollTo=ITcPnLkLuDDE. Sequences of each subunit were searched against genetic databases with msa_method = mmseqs2, pair_mode = unpaired. The default mode of sampling options was used; num_models = 5, ptm option, num_ensemble = 1, max_cycles = 3, num_samples = 1. Total 5 models were ranked according to their Predicted Local-Distance Difference Test (pLDDT) score between 0 to 100, from low to high confidence level. Septin and exocyst subunits were input in 1:1 ratio. For each of the 32 pairs of septin and exocyst subunits, the protein sequences were entered in both orders (for example, Spn1:Sec3 and Sec3:Spn1). We found that the order of input sequence affects some prediction results. So we predicted all septin-exocyst combinations in both input sequence orders. We then selected the top septin-exocyst combinations that showed interactions in both input orders. The structural figures were drawn with PyMOL version 2.0 (Schrodinger, Inc.).

We used AlphaFold3 (https://alphafoldserver.com/) to predict the structures of fission yeast exocyst complex and septin hexamer/octamer (Abramson et al., 2024). To predict the structure of whole exocyst complex, we trimmed some of the exocyst subunits to meet the 5,000-residue limit of AlphaFold3 based on the budding yeast cryo-EM structure of exocyst complex (PDB: 5YFP, 4.4 Å resolution) (Mei et al., 2018). The truncations were selected so that they do not interfere with inter-subunit interactions as well as with septin binding based on our modeling. Sequences of all subunits of the respective complexes were used as input and models were generated using the default settings. Top ranked models based on PAE, pTM, and iPTM were analyzed in PyMol. Different subunits were colored distinctly to differentiate the interface. To evaluate accessibility of residues, surface exposure of predicted interacting residues was mapped onto the corresponding residues in the final models and colored yellow to visualize distinctly.

### Co-IP and Western blotting

We carried out Co-IP and Western blotting as previously described (Laporte et al., 2011; Lee and Wu, 2012; Ye et al., 2012). Briefly, mEGFP, GFP, mYFP, or 13Myc tagged septin or exocyst subunits were expressed under native promotors in fission yeast. Cells were grown in YE5S liquid medium at 25°C for ∼48 h before harvesting and lyophilization. Lyophilized cells (200 mg) were ground into a homogeneous fine powder using pestles and mortars. IP buffer (50 mM 4-(2-hydroxyethyl)-1-piperazineethanesulfonic acid [HEPES], pH 7.5, 150 mM NaCl, 1 mM EDTA, 0.1% NP-40, 50 mM NaF, 20 mM glycerophosphate, 0.1 mM Na_3_VO_4_, 1 mM PMSF, and protease inhibitor [Roche] 1 tablet/30 ml buffer) was added according to the ratio of 10 µl : 1 mg lyophilized cell powder. 60 µl Dynabeads protein G beads (Invitrogen) were incubated with 5 µg polyclonal GFP antibody (Novus Bio) for 1 h at room temperature. After three washes with PBS and one wash with 1 ml IP buffer, the beads were incubated with cell lysate for 2 h at 4°C. After 5 washes at 4°C with 1 ml IP buffer each time, proteins were eluted by boiling with 80 µl sample buffer. The protein samples were separated with SDS-PAGE gel and detected with monoclonal anti-GFP antibody (1:1,000 dilution; 11814460001; Roche, Mannheim, Germany), monoclonal anti-Myc antibody (1:500 dilution, 9E10, Santa Cruz Biotechnology, Dallas, TX), and anti-tubulin TAT1 antibody at 1:10,000 dilution (Woods et al., 1989). Secondary antibody anti-mouse immunoglobulin G (1:5,000 dilution; A4416, Sigma-Aldrich) was detected using SuperSignal Maximum Sensitivity Substrate (Thermo Fisher Scientific) on iBright CL1500 imager (Thermo Fisher Scientific).

### Electron microscopy

Electron microscopy was performed at the Boulder Electron Microscopy Services at the University of Colorado, Boulder (Boulder, CO) as previously described (Lee et al., 2014; Wang et al., 2016). Briefly, yeast cells were grown at 25°C for ∼41 h in YE5S medium and then shifted to 36°C for 4 h before harvesting using Millipore filters. Samples were prepared using high-pressure freezing with a Wohlwend Compact 02 Freezer in the presence of 2% osmium tetroxide and 0.1% uranyl acetate in acetone. Thin sections with a thickness of 70 nm were cut and embedded in Epon-Araldite epoxy resin, which were post stained with uranyl acetate and lead citrate. Imaging of EM samples was done using a Philips CM100 transmission electron microscope (FEI, Hillsboro, OR).

### Yeast two hybrid assays

Yeast two hybrid assays were performed as described previously using X-gal overlay and β-D-galactosidase activity quantifications (Amberg et al., 2006; Paiano et al., 2019). DNA or cDNA (for genes with introns) sequences of Spn1, Spn1(aa 300-469), Spn2, Spn4, Sec5, Sec6, and Sec15 were cloned into pVP16 or pGBT9 vectors having VP16 transcription activation domain (AD) or GAL4 transcription factor DNA-binding domain (BD), respectively. Constructed plasmids were confirmed by restriction digestions and Sanger sequencing. Pairs of plasmids were then co-transformed into *S. cerevisiae* strain MAV203 (11281-011; Invitrogen) and plated on synthetic drop-out medium lacking leucine and tryptophan (SD-L-W) for selection. For X-gal overlay assay, grown colonies were re-streaked on YPD (yeast extract-peptone-dextrose) plates to grow overnight. We used 10-12 ml chloroform per plate to permeabilize cells for 10 min and then dried for 10 additional min. 0.5% agarose was prepared in 25 ml PBS (pH 7.5) and 500 µl X-gal (20 mg/ml stock in DMSO) was added after cooling. After mixing thoroughly, agarose containing X-gal was overlaid onto the colonies and incubated at 30°C. Plates were checked every 30 min for development of blue color.

Interactions were then quantified by β-D-galactosidase activity using the *o*-nitrophenyl– β-D-galactopyranoside (ONPG) assay (48712-M; Sigma Aldrich) according to the published methods (Amberg et al., 2006; Paiano et al., 2019). For interactions between Sec15 with Spn1, Spn2, and Spn4, the Amberg et al. method was used (Amberg et al., 2006). Briefly, cells were grown in SD-L-W liquid medium at 30°C overnight. 40 ml culture with OD_595_ >1 was collected and washed with 1 ml distilled water. Then cells were broken in 110 µl breaking buffer (100 mM Tris-Cl, pH 7.5, 1 mM DTT, and 20% glycerol) using glass beads on bead beater. 10 µl of the lysate was diluted with 90 µl distilled water and spun down to remove cell debris, and the supernatant was used to estimate protein concentration by Bradford assay. To the remaining 100 µl of lysate, 0.9 ml Z-buffer (100 mM sodium phosphate, pH 7.5, 10 mM KCl, and 2 mM MgSO_4_) and 0.2 ml ONPG (8 mg/1 ml Z buffer) were added and incubated at 28°C until pale yellow color developing in at least one of the samples. All the reactions were stopped by adding 0.4 ml 1 M Na_2_CO_3_. Debris were removed by centrifuging at 15,700 g for 10 min and OD_420_ was measured using 1 ml of supernatant. Time elapsed from adding ONPG to adding stop solution was recorded and activity of β-galactosidase was calculated using the formula:

β-galactosidase activity (nmol/min/mg) = OD_420_ × 1.7/ [0.0045 × protein (mg/ml) × extract volume (ml) × time (min)]

For the interaction between Spn1 and Sec6, the Painano et al. method was used (Paiano et al., 2019). Briefly, cultures were diluted to OD_595_ = 0.30 and incubated for 2 h at 30°C. For each sample, cells from 9 ml culture were collected and washed with 1 ml Z buffer and then resuspended in 0.1 ml Z buffer. Cells were broken by three freeze-thaw cycles in liquid nitrogen. 0.7 ml Z buffer with β-mercaptoethanol (27 µl β-mercaptoethanol in 9.973 ml Z buffer) and 160 µl ONPG was added to the cell lysates and incubated at 30°C until a yellow color developing in at least one of the samples. Reactions were stopped by adding 0.4 ml 1 M Na_2_CO_3_. Debris were removed by centrifuging at 15,700 g for 10 min and OD_420_ was measured using 1 ml supernatant. Time elapsed from adding ONPG to adding stop solution was recorded and β-galactosidase activity were calculated using the following formula: β-galactosidase Units = 1000 × OD_420_/ [T × V × OD_595_], where T is the elapsed time (minutes), V is the volume (ml) of culture used, and OD595 is the optical density of yeast culture.

### Statistical analysis

Data in graphs are mean ± 1 SD except where noted. The *p*-values in statistical analyses were calculated using the two-tailed Student’s *t* tests except Figure 4 (E and F), where one way ANOVA with Tukey’s post hoc test was used to quantify yeast two hybrid analyses.

## Acknowledgements

We thank Mohan Balasubramanian, Sophie Martin, Pilar Pérez, John Pringle, and Takashi Toda for fission yeast strains; Eileen O’Toole and Garry Morgan at University of Colorado, Boulder for help with electron microscopy; Anita Hopper, Steve Osmani, Dmitri Kudryashov, Elena Kudryashova, Damien Wilburn, and Emily Vais for equipment and technical support; and members of the Wu laboratory for helpful discussion and suggestions.

## Author Contributions

Performed the experiments, methodology, validation, and formal data analysis: D. Singh, Y. Liu, Y.-H. Zhu, S. Zhang, S. Naegele. Visualization, figure and table preparations: D. Singh and Y. Liu. Writing and editing manuscript: Y. Liu, D. Singh, and J.-Q. Wu. Supervision, conceptualization, investigation, review: J.-Q. Wu. Funding acquisition: Y. Liu and J.-Q. Wu.

## Additional files

### Supplementary files

Video 1. Accumulation of the septin Spn1-mEGFP and the exocyst Exo70-tdTomato to the division site.

Video 2. Dynamic localization of Exo70-tdTomato at the division site on a single-focal plane close to the cell surface.

Video 3. Dynamic localization of Exo70-tdTomato at the division site on the middle-focal plane.

Video 4. AlphaFold modeling of fission yeast exocyst complex.

Video 5. AlphaFold modeling of fission yeast septins (Spn1, 2, 3, and 4) octameric complex.

Video 6. AlphaFold modeling of septins Spn1, Spn2, Spn4 hexameric complex.

Video 7. AlphaFold modeling of septins Spn1, Spn3, Spn4 hexameric complex.

Video 8. Localization of Sec15-mEGFP in WT cells.

Video 9. The mislocalization of Sec15-mEGFP on the division plane in *spn1Δ* cells during late-stage of cytokinesis.

Video 10. The mislocalization of Sec15-mEGFP on the division plane in *spn4Δ* cells during late-stage of cytokinesis.

## Data availability

All data are available in the main text or the supplementary materials.

## Video Legends

**Video 1. Accumulation of septin Spn1-mEGFP and the exocyst marked by Exo70-tdTomato to the division site.** The cell (strain JW9170) was imaged on a single-focal plane at the cell surface every 10 s in time-lapse TIRF microscopy (Nikon Ti Microscope). DIC image shows the cell had no septa at the beginning of the movie. Scale bar, 5 μm. Display rate: 10 frames per second (fps).

**Video 2. Dynamic localization of Exo70-tdTomato at the division site on a single-focal plane close to the cell surface.** The cell (strain JW9170) was imaged without delay (500 ms exposure) in time-lapse TIRF microscopy (Nikon Ti Microscope). DIC image shows the cell had no septa at the beginning of the movie. Scale bar, 5 μm. Display rate: 10 fps.

**Video 3. Dynamic localization of Exo70-tdTomato at the division site on the middle-focal plane.** The cell (strain JW9170) was imaged without delay (500 ms exposure) in time-lapse microscopy (Nikon Ti Microscope). DIC image shows the cell had no septa at the beginning of the movie. Scale bar, 5 μm. Display rate: 5 fps.

**Video 4. The predicted 3D structural model of *S. pombe* exocyst complex by AlphaFold3, highlighting the residues that interact with septins.** Individual subunits are colored distinctly and labeled. Surface exposed residues previously predicted as putative septin-interacting sites are highlighted in yellow.

**Video 5. The predicted 3D structural model of *S. pombe* septin octameric complex by AlphaFold3, highlighting the exocyst-interacting residues.** Two subunits of each septin Spn1 to Spn4 are used to construct the octameric complex. Different subunits are colored distinctly and labeled. Surface exposed residues previously predicted as putative exocyst-interacting sites are highlighted in yellow.

**Video 6. The predicted 3D structural model of *S. pombe* septins Spn1, Spn2, and Spn4 hexameric complex by AlphaFold3, highlighting the exocyst-interacting residues.** Two subunits of each Spn1, Spn2, and Spn4 are used to construct the hexameric complex. Different subunits are colored distinctly and labeled. Surface exposed residues previously predicted as putative exocyst-interacting sites are highlighted in yellow.

**Video 7. The predicted 3D structural model of *S. pombe* septins Spn1, Spn3, and Spn4 hexameric complex by AlphaFold3, highlighting the exocyst-interacting residues.** Two subunits of each Spn1, Spn3, and Spn4 are used to construct the hexameric complex. Different subunits are colored distinctly and labeled. Surface exposed residues previously predicted as putative exocyst-interacting sites are highlighted in yellow.

**Video 8. Localization of Sec15 as a ring at the rim of the division plane during cytokinesis in WT cells.** Sec15-mEGFP cells (strain JW9726) were imaged at 3 min interval for 2 h in time-lapse confocal microscopy (UltraVIEW Vox CSUX1; PerkinElmer). 3D projections of fluorescence images from 14 slices spaced at 0.4 µm at each time point are shown. Display rate: 50 fps.

**Video 9. Mislocalization of Sec15 as a disk on division site in *spn1Δ* cells.**

*sec15-mEGFP spn1Δ* cells (strain JW9852) were imaged at 3 min interval for 3 h in time-lapse confocal microscopy (UltraVIEW Vox CSUX1; PerkinElmer). 3D projections of fluorescence images from 14 slices spaced at 0.4 µm at each time point are shown. Display rate: 50 fps.

**Video 10. Mislocalization of Sec15 as a disk on division site in *spn4Δ* cells.**

*sec15-mEGFP spn4Δ* cells (strain JW9853) were imaged at 3 min interval for 3 h in time-lapse confocal microscopy (UltraVIEW Vox CSUX1; PerkinElmer). 3D projections of fluorescence images from 14 slices spaced at 0.4 µm at each time point are shown. Display rate: 50 fps.

---

**Figure S1.**
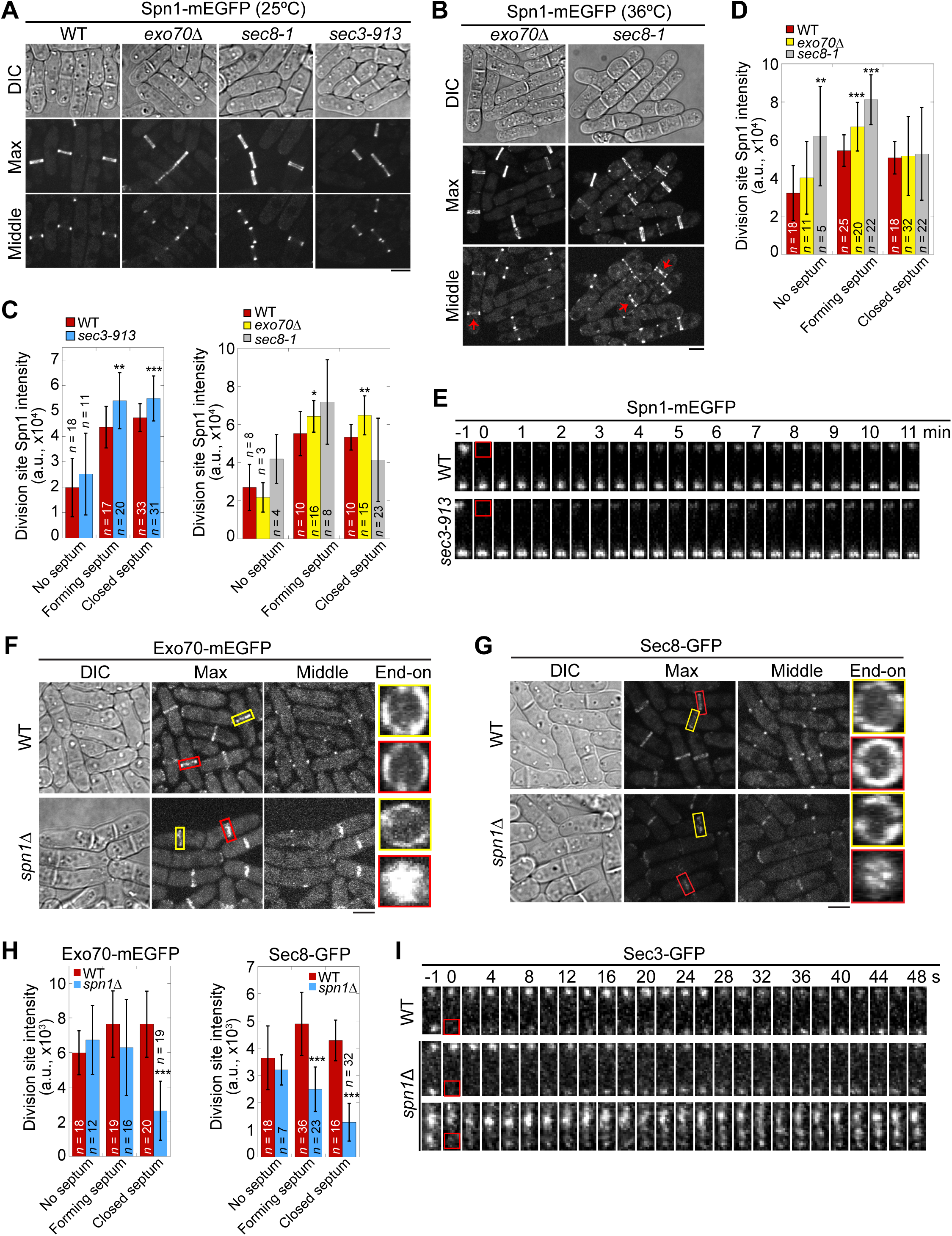
Localization and division site levels of septin and exocyst subunits in mutants; and FRAP analyses of Spn1 and Sec3. **(A and B)** Localization of Spn1 in WT and exocyst mutants at 25°C (A) and 4 h at 36°C (B). Arrows indicate cells with mislocalized Spn1 at the center of the division plane. **(C and D)** Quantifications of Spn1 intensities at the division site in WT and exocyst mutants at 25°C (C) and 4 h at 36°C (D). No septum: cells with Spn1 signal at the division site but no septum is visible under DIC; forming septum: septum with a visible gap in the middle; closed septum: no visible gap in the septum. *, P < 0.05; **, P < 0.01; ***, P < 0.001. **(E)** FRAP analyses of Spn1 at the division site in WT and *sec3-913* cells grown at 36°C for 4 h. Time-lapse images show recovery of Spn1 signals over time. Red box marks the region photobleached at time 0. **(F and G)** Localization of Exo70 (F) and Sec8 (G) in WT and *spn1*Δ cells. Yellow boxes, cells without a septum; Red boxes, cells with a closed septum. **(H)** Quantifications of Exo70 (left) and Sec8 (right) intensities at the division site in WT and *spn1*Δ cells. ***, P < 0.001. **(I)** FRAP analyses of Sec3 at the division site in WT and *spn1*Δ cells. Red box marks the region photobleached at time 0. Bars, 5 μm.

**Figure S2.**
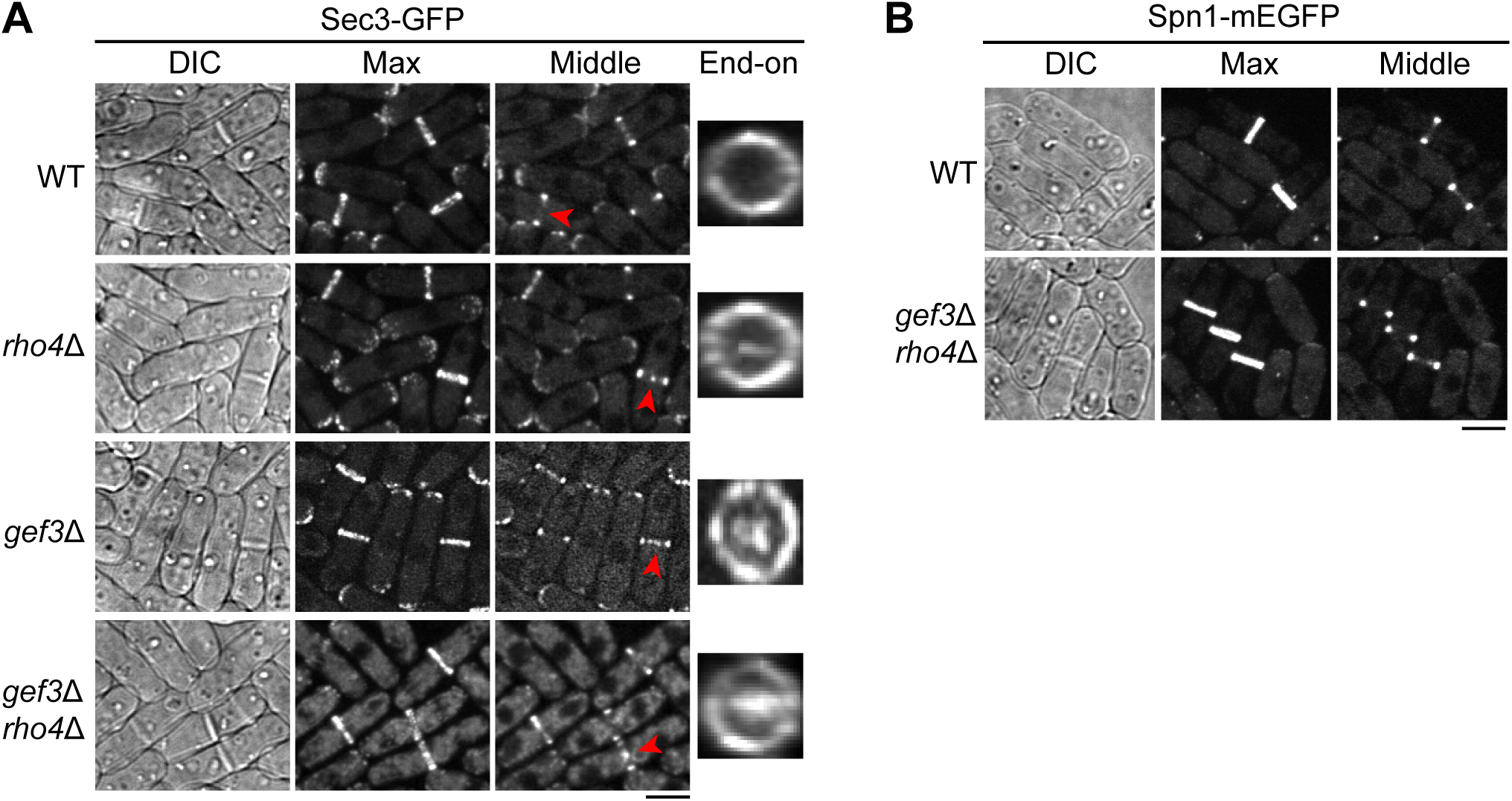
Sec3 and Spn1 localization in *gef3*, *rho4*, or *gef3 rho4* mutants. (A) Sec3 localization in WT, *rho4*Δ, *gef3*Δ, and *gef3*Δ *rho4*Δ cells. Arrowheads mark examples of the cells with mislocalized Sec3 at the center of the division plane in mutant but not WT cells. End-on views of the division plane of cells with a closed septum are shown on the last column. (B) Spn1 localization in WT and *gef3*Δ *rho4*Δ cells. Bars, 5 μm.

**Figure S3.**
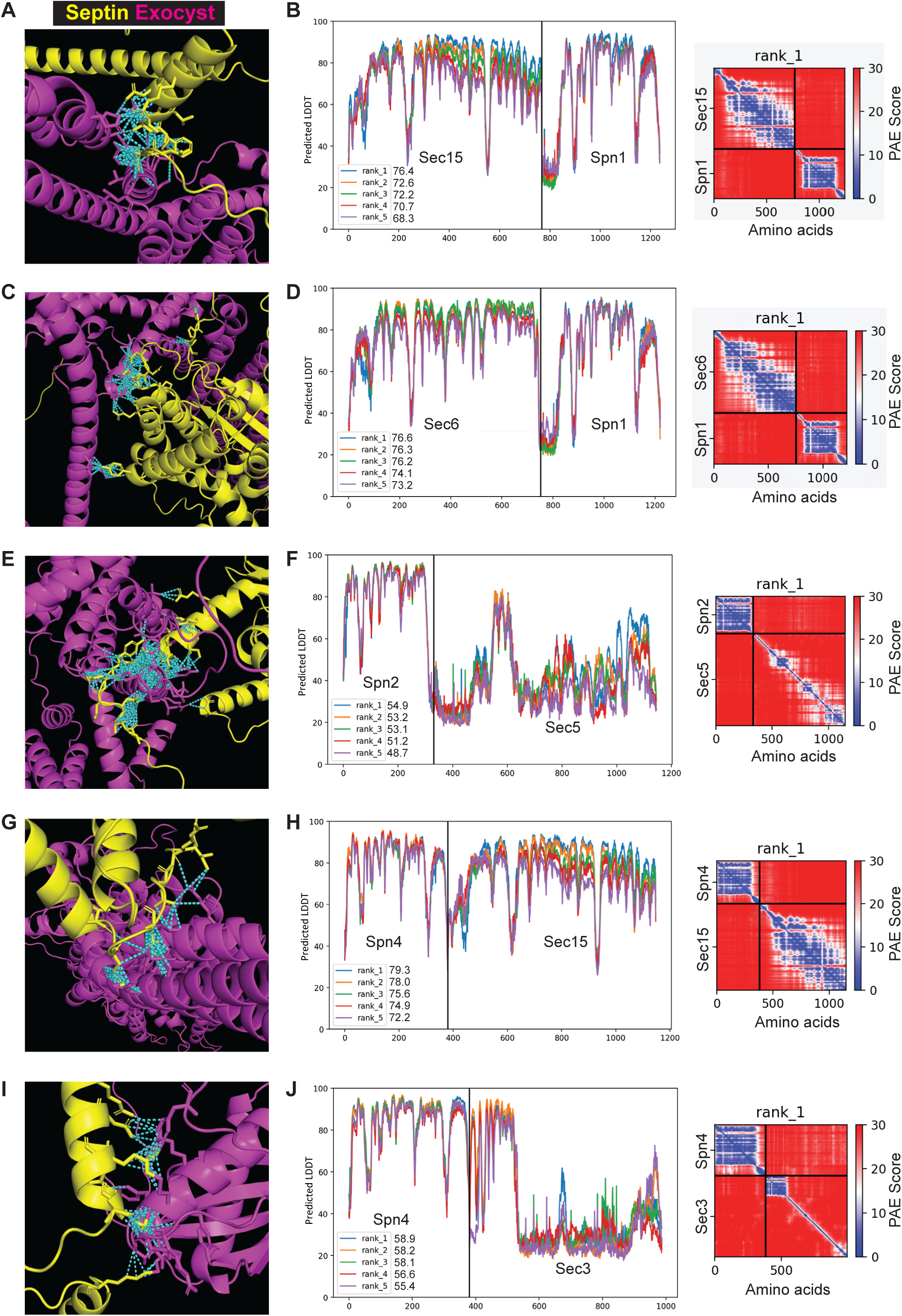
The 3D structural models of septin-exocyst interactions generated by AlphaFold. **(A, C, E, G, I)** Rank 1 model of AlphaFold2_advanced predicted interaction between Sec15 and Spn1 (A, pTM score = 0.47), Sec6 and Spn1 (C, pTM = 0.45), Spn2 and Sec5 (E, pTM = 0.37), Spn4 and Sec15 (G, pTM = 0.48), and Spn4 and Sec3 (I, pTM = 0.43). Septin subunits are colored in yellow and the exocyst in magenta, contacts between interface residues with distance < 4 Å are colored in cyan**. (B, D, F, H, J)** pLDDT scores of five predicted models and the PAE plot of rank1 model for Sec15 and Spn1 (B), Sec6 and Spn1 (D), Spn2 and Sec5 (F), Spn4 and Sec15 (H), and Spn4 and Sec3 (J).

**Figure S4.**
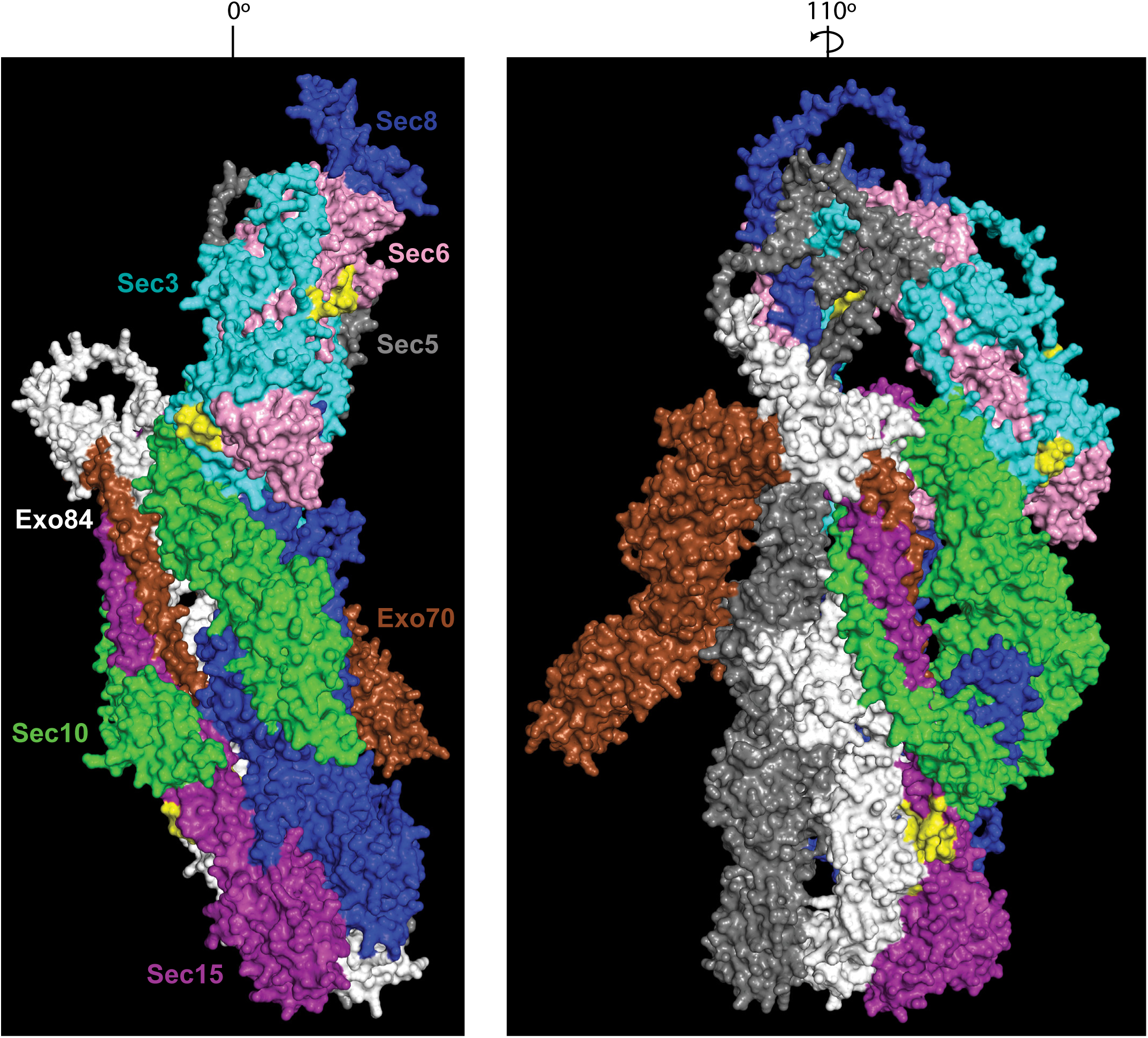
The predicted 3D structural model of *S. pombe* exocyst complex by AlphaFold3, highlighting the residues that interact with septins (also see Video 4). Individual subunits are colored distinctly and labeled. Surface exposed residues previously identified as putative septin-interacting sites are highlighted in yellow. The model demonstrates that ∼84% predicted septin-interacting residues are accessible on the outer surface of the assembled complex. As AlphaFold3 limits to fit the whole complex in 5000 tokens, full length Sec5, Exo70, Ex84 were used, but only amino acids 1-500 for Sec3, 1-546 for Sec6, 1-865 for Sec8, 1-620 for Sec10, and 1-396 for Sec15 were used. These truncations were selected based on the budding yeast exocyst cryo-EM structure (PDB: 5YFP), which shows that these regions are sufficient for stable inter-subunit interactions and are unlikely to interfere with septin binding based on our modeling.

**Figure S5.**
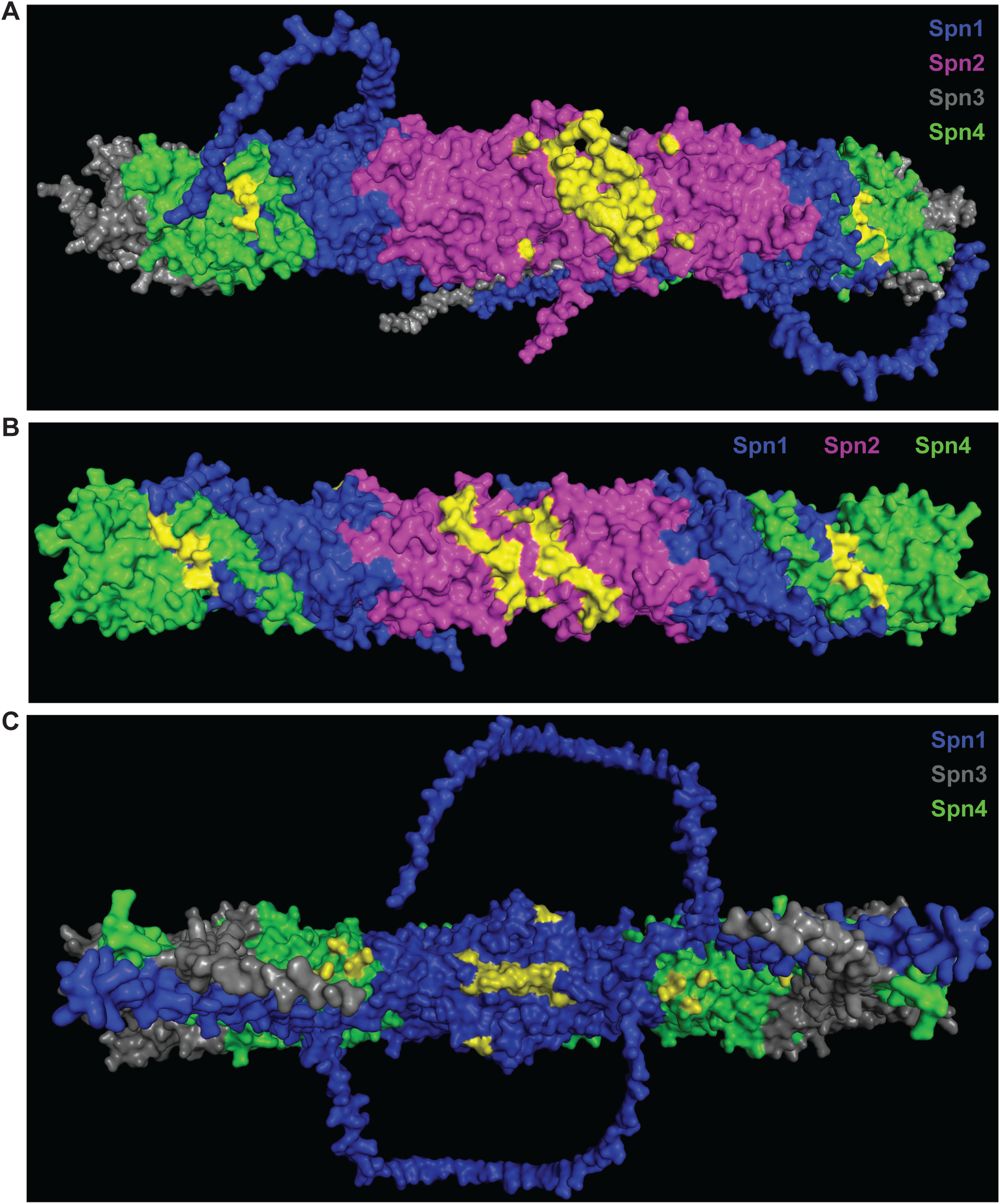
The predicted 3D structural models of *S. pombe* septin complexes by AlphaFold3, highlighting the exocyst-interacting residues (Videos 5-7). (A-C) Two subunits of each septin were used to construct the octameric or hexameric complex. Different subunits are colored distinctly and labeled. Surface exposed residues previously identified as putative exocyst-interacting sites are highlighted in yellow. (A) The octameric model of Spn1 to Spn4 demonstrates that ∼96% predicted exocyst-interacting residues are accessible on the outer surface of the assembled complex. (B) The hexameric complex of two subunits of each Spn1, Spn2, and Spn4 shows 92%, (C) of each Spn1, Spn3 and Spn4 shows 86% exocyst-interacting residues are available on outer surface of assembled complex.

**Figure S6.**
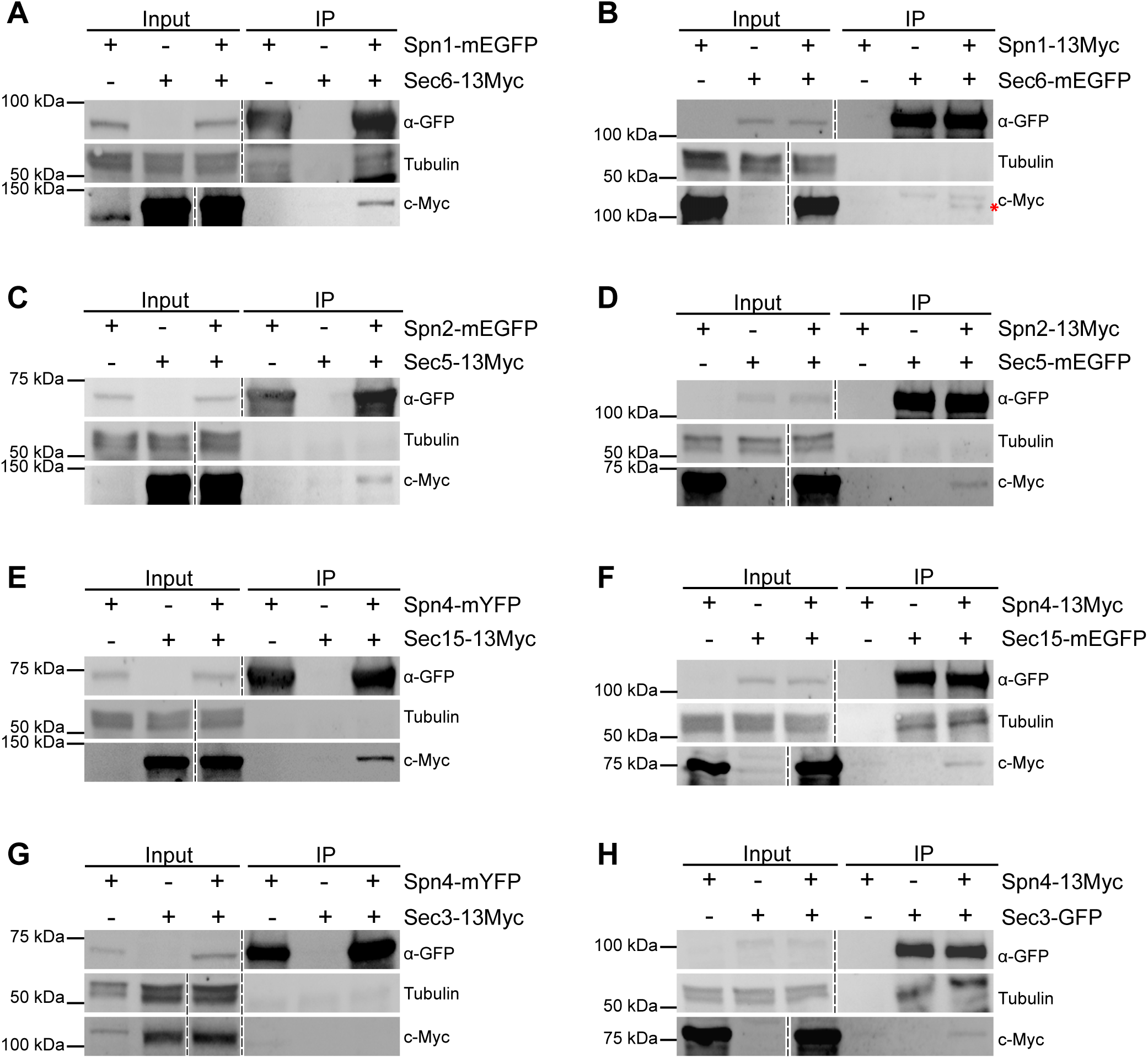
Septins and the exocyst interact physically. Reciprocal co-immunoprecipitation between Spn1 with Sec6 (A, B); Spn2 with Sec5 (C, D); Spn4 with Sec15 (E, F); and Spn4 with Sec3 (G, H). Septin or exocyst subunits tagged with mEGFP, GFP, mYFP, or 13Myc were immunoprecipitated, separated on SDS-PAGE, and incubated with appropriate antibodies. Tubulin was used as a loading control. Asterisk (*) in B marks Spn1-13Myc. The dashed vertical lines mark the positions of protein ladders which were excised out.

**Supplemental Table 1.** *S. pombe* strains used in this study.

| Strain | Genotype | Figure; Video; Table; (Reference) |
| --- | --- | --- |
| JW8692 | *sec3-tdTomato-hphMX6 spn1-mEGFP-kanMX6 sad1-mRFP1-kanMX6 ade6-M210 leu1-32 ura4-D18* | Figure 1, A, B, D and E |
| JW9170 | *spn1-mEGFP-kanMX6 exo70-tdTomato-natMX6 ade6-M210 leu1-32 ura4-D18* | Figure 1C; Videos 1-3 |
| JW1113 | *h<sup>-</sup> spn1-mEGFP-kanMX6 sad1-mRFP1-kanMX6 ade6-M210 leu1-32 ura4-D18* | Figure 1F |
| JW1100 | *spn1-mEGFP-kanMX6 ade6-M210 leu1-32 ura4-D18* | Figures 1, H and I; 4C; S2B; S6A |
| JW8928 | *spn1-mEGFP-kanMX6 sec3-913-hphMX6 ade6 leu1-32 ura4-D18* | Figures 1, F, H and I; S1, A, C and E |
| JW8848 | *sec8-1 spn1-mEGFP-kanMX6 sad1-mRFP1-kanMX6 rng8-tdTomato-kanMX6 ade6-M210 leu1-32 ura4-D18* | Figure 1G |
| IJ612 | *h<sup>+</sup> sec3-GFP-kanMX6 ade6-M216 leu1-32 ura4-D18* | Figures 2, A-C; S1I; S2A; (Jourdain et al., 2012) |
| JW7322 | *spn1-Δ2::kanMX6 sec3-GFP-kanMX6 ade6-M216 leu1-32 ura4-D18* | Figures 2, A-C; S1I |
| JW7061 | *h<sup>-</sup> sec8-GFP-ura4<sup>+</sup> rlc1-tdTomato-natMX6 ade6<sup>?</sup> leu1-32 ura4-D18* | Figures 2, D and E; S1, G and H |
| JW8295 | *sec8-GFP-ura4<sup>+</sup> rlc1-tdTomato-natMX6 spn1-Δ2::kanMX6 leu1-32 ura4-D18* | Figures 2, D and E; S1, G and H |
| JW9737 | *h<sup>+</sup> spn2-mEGFP-kanMX6 ade6-M210 ura4-D18 leu1-32* | Figure 4A |
| JW9731 | *h<sup>-</sup> sec15-13Myc-kanMX6 ade6-210 ura4-D18 leu1-32* | Figures 4, A and C; S6E |
| JW9757 | *spn2-mEGFP-kanMX6 sec15-13Myc-kanMX6 ade6-M210 ura4-D18 leu1-32* | Figure 4A |
| JW9756 | *h<sup>+</sup> spn2-13Myc-kanMX6 ade6-M210 leu1-32 ura4-D18* | Figure 4B |
| JW9726 | *h<sup>-</sup> sec15-mEGFP-kanMX6 ade6-210 ura4-D18 leu1-32* | Figure 4, B and D; Video 8 |
| JW9771 | *spn2-13Myc-kanMX6 sec15-mEGFP-kanMX6 ade6-210 ura4-D18 leu1-32* | Figure 4B |
| JW9744 | *h<sup>+</sup> spn1-mEGFP-kanMX6 sec15-13Myc-kanMX6 ade6-M210 leu1-32 ura4-D18* | Figure 4C |
| JW9733 | *h<sup>-</sup> spn1-13Myc-hphMX6 sec15-mEGFP-kanMX6 ade6-M210 ura4-D18 leu1-32* | Figure 4D |
| JW8596 | *h<sup>+</sup> spn1-13Myc-hphMX6 ade6-M210 leu1-32 ura4-D18* | Figures 4D; S6B |
| JW9789 | *h<sup>+</sup> spn2-Δ1::hphMX6 sec15-mEGFP-kanMX6 ade6-210 ura4-D18 leu1-32* | Figure 5, A and B |
| JW9759 | *h<sup>+</sup> sec15-mEGFP-kanMX6 ade6-210 ura4-D18 leu1-32* | Figures 5, A and B; S6F |
| JW9852 | *sec15-mEGFP-kanMX6 spn1-Δ2::kanMX6 ade6-M210? ura4-D18 leu1-32* | Figure 5, A and B; Video 9 |
| JW9853 | *sec15-mEGFP-kanMX6 spn4-Δ2::hphMX6 ade6-M210 ura4-D18 leu1-32* | Figure 5, A and B; Video 10 |
| JW9804 | *spn2-Δ1::hphMX6 sec5-mEGFP-kanMX6 ade6-210 ura4-D18 leu1-32* | Figure 5, C and D |
| JW9791 | *h<sup>+</sup> sec5-mEGFP-kanMX6 ade6-M210 ura4-D18 leu1-32* | Figure 5, C and D |
| JW81 | *h<sup>-</sup> ade6-210 ura4-D18 leu1-32* | Figures 6, A and B; 7B |
| JW289 | *h<sup>+</sup> spn1-Δ2::kanMX6 leu1-32 ura4-D18* | Figures 6, A and B; 7B; Table 1 |
| MBY887 | *h<sup>+</sup> sec8-1 ura4-D18 leu1-32* | Figures 6, A and B; 7B; (Wang et al., 2002) |
| JW7130 | *h<sup>-</sup> kanMX6-Pypt3-mEGFP-ypt3 ade6-210 leu1-32 ura4-D18* | Figure 6C |
| JW7354 | *spn1-Δ2::kanMX6 kanMX6-Pypt3-mEGFP-ypt3 leu1-32 ura4-D18* | Figure 6C |
| JW6548 | *h<sup>+</sup> GFP-syb1-kanMX6 rlc1-tdTomato-natMX6 ade6 leu1-32 ura4-D18* | Figure 6D |
| JW7385 | *spn1-Δ2::kanMX6 GFP-syb1-kanMX6 rlc1-tdTomato-natMX6 leu1-32 ura4-D18* | Figure 6D |
| JW5249 | *GFP-bgs1-leu1<sup>+</sup> bgs1Δ::ura4<sup>+</sup> rlc1-tdTomato-natMX6 ade6-M210 leu1-32 ura4-D18* | Figure 7A |
| JW7264 | *GFP-bgs1-leu1<sup>+</sup> bgs1Δ::ura4<sup>+</sup> rlc1-tdTomato-natMX6 spn1-Δ2::kanMX6 ade6 ura4-D18* | Figure 7A |
| PPG37.23 | *h<sup>-</sup> eng1-GFP-kan<sup>R</sup> leu1-32 ura4-D18* | Figure 7C (Santos et al., 2005) |
| JW9057 | *eng1-GFP-kan<sup>R</sup> spn1-Δ2::kanMX6 leu1-32 ura4-D18* | Figure 7C |
| JW1113 | *h<sup>-</sup> spn1-mEGFP-kanMX6 sad1-mRFP1-kanMX6 ade6-M210 leu1-32 ura4-D18* | Figure S1, A, C-E; |
| JW8829 | *exo70Δ::kanMX4 spn1-mEGFP-kanMX6 sad1-mRFP1-kanMX6 ade6 leu1-32 ura4-D18* | Figure S1, A-D |
| JW8830 | *sec8-1 spn1-mEGFP-kanMX6 sad1-mRFP1-kanMX6 ade6-M210 leu1-32 ura4-D18* | Figure S1, A-D |
| JW8929 | *h<sup>-</sup> exo70-mEGFP-kanMX6 ade6-M210 ura4-D18 leu1-32* | Figure S1, F and H |
| JW8960 | *exo70-mEGFP-kanMX6 spn1-Δ2::kanMX6 ade6-M210 leu1-32 ura4-D18* | Figure S1, F and H |
| JW8938 | *rho4Δ::kanMX6 sec3-GFP-kanMX6 leu1-32 ura4-D18* | Figure S2A |
| JW8955 | *gef3Δ::hphMX6 sec3-GFP-kanMX6 ade6-M210 leu1-32 ura4-D18* | Figure S2A |
| JW8959 | *gef3Δ::hphMX6 rho4Δ::kanMX4 sec3-GFP-kanMX6 ade6? leu1-32? ura4-D18* | Figure S2A |
| JW9058 | *gef3Δ::hphMX6 rho4Δ::kanMX4 spn1-mEGFP-kanMX6 ade6 leu1-32 ura4-D18* | Figure S2B |
| JW9765 | *h<sup>-</sup> sec6-13Myc-kanMX6 ade6-210 ura4-D18 leu1-32* | Figure S6A |
| JW9774 | *spn1-mEGFP-kanMX6 sec6-13Myc-kanMX6 ade6-M210 leu1-32 ura4-D18* | Figure S6A |
| JW9766 | *h<sup>-</sup> sec6-mEGFP-kanMX6 ade6-210 ura4-D18 leu1-32* | Figure S6B |
| JW9832 | *spn1-13Myc-hphMX6 sec6-mEGFP-kanMX6 ade6-210 ura4-D18 leu1-32* | Figure S6B |
| JW8778 | *h<sup>-</sup> spn2-mEGFP-kanMX6 ade6-M210 leu1-32 ura4-D18* | Figure S6C |
| JW9755 | *h<sup>+</sup> sec5-13Myc-kanMX6 ade6-M210 leu1-32 ura4-D18* | Figure S6C |
| JW9772 | *sec5-13Myc-kanMX6 spn2-mEGFP-kanMX6 ade6-210 ura4-D18 leu1-32* | Figure S6C |
| JW9756 | *h<sup>+</sup> spn2-13Myc-kanMX6 ade6-M210 leu1-32 ura4-D18* | Figure S6D |
| JW9738 | *h<sup>-</sup> sec5-mEGFP-kanMX6 ade6-M210 ura4-D18 leu1-32* | Figure S6D |
| JW9775 | *spn2-13Myc-kanMX6 sec5-mEGFP-kanMX6 ade6-M210 leu1-32 ura4-D18* | Figure S6D |
| JW1171 | *h<sup>+</sup> spn4-mYFP-kanMX6 ade6-M210 leu1-32 ura4-D18* | Figure S6, E and G |
| JW9829 | *sec15-13Myc-kanMX6 spn4-mYFP-kanMX6 ade6-M210 ura4-D18 leu1-32* | Figure S6E |
| JW9854 | *spn4-13Myc-kanMX6 sec15-mEGFP-kanMX6 ade6-210 ura4-D18 leu1-32* | Figure S6F |
| JW9768 | *h<sup>-</sup> spn4-13Myc-kanMX6 ade6-210 ura4-D18 leu1-32* | Figure S6, F and H |
| JW9139 | *h<sup>-</sup> sec3-13Myc-natMX6 ade6-210 leu1-32 ura4-D18* | Figure S6G |
| JW9711 | *spn4-mYFP-kanMX6 sec3-13Myc-natMX6 ade6-M210 leu1-32 ura4-D18* | Figure S6G |
| JW7300 | *h<sup>+</sup> sec3-GFP-kanMX6 ade6-M210 leu1-32 ura4-D18* | Figure S6H |
| JW9788 | *spn4-13Myc-kanMX6 sec3-GFP-kanMX6 ade6-210 leu1-32 ura4-D18* | Figure S6H |
| JW7035 | *h<sup>-</sup> trs120-M1-his5<sup>+</sup>-kanMX6 his5Δ ade6-M210 leu1-32 ura4* | Tables 1 and 2 |
| JW8821 | *trs120-M1-his5<sup>+</sup>-kanMX6 spn1-Δ2::kanMX6 leu1-32 ura4* | Tables 1 and 2 |
| JW7036 | *h<sup>-</sup> trs120-ts1-his5<sup>+</sup>-kanMX6 his5Δ ade6-M210 leu1-32 ura4* | Tables 1 and 2 |
| JW8822 | *trs120-ts1-his5<sup>+</sup>-kanMX6 spn1-Δ2::kanMX6 leu1-32 ura4* | Tables 1 and 2 |
| JW290 | *h<sup>-</sup> spn1-Δ2::kanMX6 his3-27 ura4-D18* | Tables 1 and 2 |
| MBY887 | *h<sup>+</sup> sec8-1 ura4-D18 leu1-32* | Tables 1 and 2 |
| JW8796 | *spn1-Δ2::kanMX6 sec8-1 ura4-D18* | Tables 1 and 2 |
| JW2716 | *h<sup>+</sup> exo70Δ::kanMX4 ade6 leu1-32 ura4-D18* | Tables 1 and 2 |
| JW8797 | *spn1-Δ2::kanMX6 exo70Δ::kanMX4 his3-27 ade6 ura4-D18* | Tables 1 and 2 |
| IJ1032 | *h<sup>-</sup> sec3-916-hphMX6 ade6-M216 leu1-32 ura4-D18* | Tables 1 and 2;<br>(Jourdain et al., 2012) |
| JW8787 | *spn1-Δ2::kanMX6 sec3-916-hphMX6 ade6-M216 leu1-32 ura4-D18* | Tables 1 and 2 |
| IJ767 | *h<sup>-</sup> sec3-913-hphMX6 ade6-M216 leu1-32 ura4-D18* | Tables 1 and 2;<br>(Jourdain et al., 2012) |
| JW8783 | *spn1-Δ2::kanMX6 sec3-913-hphMX6 ade6-M216 leu1-32 ura4-D18* | Tables 1 and 2 |
| JW8588 | *spn2-Δ1::ura4<sup>+</sup> ade6 leu1-32 ura4-D18* | Tables 1 and 2 |
| JW8782 | *spn2-Δ1::ura4<sup>+</sup> sec3-913-hphMX6 ade6 leu1-32 ura4-D18* | Tables 1 and 2 |
| JW8789 | *spn2-Δ1::ura4<sup>+</sup> sec3-916-hphMX6 ade6 leu1-32 ura4-D18* | Tables 1 and 2 |
| JW8590 | *h<sup>+</sup> spn3-Δ2::kanMX6 ade6-M210 leu1-32 ura4-D18* | Tables 1 and 2 |
| JW8790 | *spn3-Δ2::kanMX6 sec3-913-hphMX6 ade6-M21X leu1-32 ura4-D18* | Tables 1 and 2 |
| JW8785 | *spn3-Δ2::kanMX6 sec3-916-hphMX6 ade6-M21X leu1-32 ura4-D18* | Tables 1 and 2 |
| JW293 | *h<sup>-</sup> spn4-Δ2::kanMX6 ura4-D18* | Tables 1 and 2 |
| JW295 | *h<sup>+</sup> spn4-Δ2::kanMX6 leu1-32 ura4-D18* | Tables 1 and 2 |
| JW8784 | *spn4-Δ2::kanMX6 sec3-913-hphMX6 ade6-M216 leu1-32 ura4-D18* | Tables 1 and 2 |
| JW8788 | *spn4-Δ2::kanMX6 sec3-916-hphMX6 leu1-32 ura4-D18* | Tables 1 and 2 |
| JW8799 | *spn4-Δ2::kanMX6 sec8-1 leu1-32 ura4-D18* | Tables 1 and 2 |

## References

1. Abramson J, Adler J, Dunger J, Evans R, Green T, Pritzel A, Ronneberger O, Willmore L, Ballard AJ, Bambrick J, Bodenstein SW, Evans DA, Hung CC, O’Neill M, Reiman D, Tunyasuvunakool K, Wu Z, Žemgulytė A, Arvaniti E, Beattie C, et al. 2024. Accurate structure prediction of biomolecular interactions with AlphaFold 3. Nature 630:493–500.

2. Ageta-Ishihara N, Kinoshita M. 2021. Developmental and postdevelopmental roles of septins in the brain. Neurosci Res 170:6–12.

3. Ahmed SM, Nishida-Fukuda H, Li Y, McDonald WH, Gradinaru CC, Macara IG. 2018. Exocyst dynamics during vesicle tethering and fusion. Nat Commun 9:5140.

4. Amberg DC, Burke DJ, Strathern JN. 2006. Assay of β-galactosidase in yeast: assay of crude extracts. Cold Spring Harb Protoc 2006:pdb. prot4157.

5. An H, Morrell JL, Jennings JL, Link AJ, Gould KL. 2004. Requirements of fission yeast septins for complex formation, localization, and function. Mol Biol Cell 15:5551–5564.

6. Bähler J, Wu J-Q, Longtine MS, Shah NG, McKenzie A, III, Steever AB, Wach A, Philippsen P, Pringle JR. 1998. Heterologous modules for efficient and versatile PCR-based gene targeting in *Schizosaccharomyces pombe*. Yeast 14:943–951.

7. Baladrón V, Ufano S, Dueñas E, Martín-Cuadrado AB, del Rey F, Vázquez de Aldana CR. 2002. Eng1p, an endo-1,3-β-glucanase localized at the daughter side of the septum, is involved in cell separation in *Saccharomyces cerevisiae*. Eukaryot Cell 1:774–786.

8. Beites CL, Xie H, Bowser R, Trimble WS. 1999. The septin CDCrel-1 binds syntaxin and inhibits exocytosis. Nat Neurosci 2:434–439.

9. Bendezú FO, Martin SG. 2011. Actin cables and the exocyst form two independent morphogenesis pathways in the fission yeast. Mol Biol Cell 22:44–53.

10. Bendezú FO, Vincenzetti V, Martin SG. 2012. Fission yeast Sec3 and Exo70 are transported on actin cables and localize the exocyst complex to cell poles. PLoS One 7:e40248.

11. Berlin A, Paoletti A, Chang F. 2003. Mid2p stabilizes septin rings during cytokinesis in fission yeast. J Cell Biol 160:1083–1092.

12. Bertin A, McMurray MA, Grob P, Park SS, Garcia G, III, Patanwala I, Ng HL, Alber T, Thorner J, Nogales E. 2008. *Saccharomyces cerevisiae* septins: supramolecular organization of heterooligomers and the mechanism of filament assembly. Proc Natl Acad Sci U S A 105:8274–8279.

13. Bertin A, McMurray MA, Thai L, Garcia G, III, Votin V, Grob P, Allyn T, Thorner J, Nogales E. 2010. Phosphatidylinositol-4,5-bisphosphate promotes budding yeast septin filament assembly and organization. J Mol Biol 404:711–731.

14. Bi E, Maddox P, Lew DJ, Salmon ED, McMillan JN, Yeh E, Pringle JR. 1998. Involvement of an actomyosin contractile ring in *Saccharomyces cerevisiae* cytokinesis. J. Cell Biol. 142:1301–1312.

15. Boyd C, Hughes T, Pypaert M, Novick P. 2004. Vesicles carry most exocyst subunits to exocytic sites marked by the remaining two subunits, Sec3p and Exo70p. J Cell Biol 167:889–901.

16. Bridges AA, Gladfelter AS. 2016. In vitro reconstitution of septin assemblies on supported lipid bilayers. Methods Cell Biol 136:57–71.

17. Bridges AA, Zhang H, Mehta SB, Occhipinti P, Tani T, Gladfelter AS. 2014. Septin assemblies form by diffusion-driven annealing on membranes. Proc Natl Acad Sci U S A 111:2146–2151.

18. Cannon KS, Woods BL, Crutchley JM, Gladfelter AS. 2019. An amphipathic helix enables septins to sense micrometer-scale membrane curvature. J Cell Biol 218:1128–1137.

19. Cannon KS, Woods BL, Gladfelter AS. 2017. The unsolved problem of how cells sense micron-scale curvature. Trends Biochem Sci 42:961–976.

20. Cao L, Ding X, Yu W, Yang X, Shen S, Yu L. 2007. Phylogenetic and evolutionary analysis of the septin protein family in metazoan. FEBS Lett 581:5526–5532.

21. Carim SC, Hickson GRX. 2023. The Rho1 GTPase controls anillo-septin assembly to facilitate contractile ring closure during cytokinesis. iScience 26:106903.

22. Caudron F, Barral Y. 2009. Septins and the lateral compartmentalization of eukaryotic membranes. Dev Cell 16:493–506.

23. Coffman VC, Nile AH, Lee I-J, Liu H, Wu J-Q. 2009. Roles of formin nodes and myosin motor activity in Mid1p-dependent contractile-ring assembly during fission yeast cytokinesis. Mol Biol Cell 20:5195–5210.

24. Coffman VC, Wu P, Parthun MR, Wu J-Q. 2011. CENP-A exceeds microtubule attachment sites in centromere clusters of both budding and fission yeast. J Cell Biol 195:563–572.

25. Cortes JC, Ishiguro J, Duran A, Ribas JC. 2002. Localization of the (1,3)β-D-glucan synthase catalytic subunit homologue Bgs1p/Cps1p from fission yeast suggests that it is involved in septation, polarized growth, mating, spore wall formation and spore germination. J Cell Sci 115:4081–4096.

26. Dagdas YF, Yoshino K, Dagdas G, Ryder LS, Bielska E, Steinberg G, Talbot NJ. 2012. Septin-mediated plant cell invasion by the rice blast fungus, *Magnaporthe oryzae*. Science 336:1590–1595.

27. Davidson R, Laporte D, Wu J-Q. 2015. Regulation of Rho-GEF Rgf3 by the arrestin Art1 in fission yeast cytokinesis. Mol Biol Cell 26:453–466.

28. Davidson R, Liu Y, Gerien KS, Wu J-Q. 2016. Real-time visualization and quantification of contractile ring proteins in single living cells. Methods Mol Biol 1369:9–23.

29. DeMarini DJ, Adams AE, Fares H, De Virgilio C, Valle G, Chuang JS, Pringle JR. 1997. A septin-based hierarchy of proteins required for localized deposition of chitin in the *Saccharomyces cerevisiae* cell wall. J Cell Biol 139:75–93.

30. Dobbelaere J, Barral Y. 2004. Spatial coordination of cytokinetic events by compartmentalization of the cell cortex. Science 305:393–396.

31. Dobbelaere J, Gentry MS, Hallberg RL, Barral Y. 2003. Phosphorylation-dependent regulation of septin dynamics during the cell cycle. Dev Cell 4:345–357.

32. Dolat L, Hu Q, Spiliotis ET. 2014. Septin functions in organ system physiology and pathology.Biol Chem 395:123–141.

33. Du Y, Mpina MH, Birch PR, Bouwmeester K, Govers F. 2015. Phytophthora infestans RXLR effector AVR1 interacts with exocyst component Sec5 to manipulate plant immunity. Plant Physiol 169:1975–1990.

34. Fang X, Luo J, Nishihama R, Wloka C, Dravis C, Travaglia M, Iwase M, Vallen EA, Bi E. 2010. Biphasic targeting and cleavage furrow ingression directed by the tail of a myosin II. J Cell Biol 191:1333–1350.

35. Feng S, Knödler A, Ren J, Zhang J, Zhang X, Hong Y, Huang S, Peränen J, Guo W. 2012. A Rab8 guanine nucleotide exchange factor-effector interaction network regulates primary ciliogenesis. J Biol Chem 287:15602–15609.

36. Finger FP, Hughes TE, Novick P. 1998. Sec3p is a spatial landmark for polarized secretion in budding yeast. Cell 92:559–571.

37. Finnigan GC, Booth EA, Duvalyan A, Liao EN, Thorner J. 2015. The carboxy-terminal tails of septins Cdc11 and Shs1 recruit myosin-II binding factor Bni5 to the bud neck in *Saccharomyces cerevisiae*. Genetics 200:843–862.

38. Ford SK, Pringle JR. 1991. Cellular morphogenesis in the *Saccharomyces cerevisiae* cell cycle: localization of the CDC11 gene product and the timing of events at the budding site. Dev Genet 12:281–292.

39. Frazier JA, Wong ML, Longtine MS, Pringle JR, Mann M, Mitchison TJ, Field C. 1998. Polymerization of purified yeast septins: evidence that organized filament arrays may not be required for septin function. J Cell Biol 143:737–749.

40. Fukai S, Matern HT, Jagath JR, Scheller RH, Brunger AT. 2003. Structural basis of the interaction between RalA and Sec5, a subunit of the sec6/8 complex. EMBO J 22:3267–3278.

41. Ganesan SJ, Feyder MJ, Chemmama IE, Fang F, Rout MP, Chait BT, Shi Y, Munson M, Sali A. 2020. Integrative structure and function of the yeast exocyst complex. Protein Sci 29:1486–1501.

42. Garcia G, III, Bertin A, Li Z, Song Y, McMurray MA, Thorner J, Nogales E. 2011. Subunit-dependent modulation of septin assembly: budding yeast septin Shs1 promotes ring and gauze formation. J Cell Biol 195:993–1004.

43. Gladfelter AS, Pringle JR, Lew DJ. 2001. The septin cortex at the yeast mother-bud neck. Curr Opin Microbiol 4:681–689.

44. Guo PP, Yong JY, Wang YM, Li CR. 2016. Sec15 links bud site selection to polarised cell growth and exocytosis in *Candida albicans*. Sci Rep 6:26464.

45. Guo W, Roth D, Walch-Solimena C, Novick P. 1999. The exocyst is an effector for Sec4p, targeting secretory vesicles to sites of exocytosis. EMBO J 18:1071–1080.

46. Guo W, Tamanoi F, Novick P. 2001. Spatial regulation of the exocyst complex by Rho1 GTPase. Nat Cell Biol 3:353–360.

47. Gupta YK, Dagdas YF, Martinez-Rocha AL, Kershaw MJ, Littlejohn GR, Ryder LS, Sklenar J, Menke F, Talbot NJ. 2015. Septin-dependent assembly of the exocyst is essential for plant infection by *Magnaporthe oryzae*. Plant Cell 27:3277–3289.

48. Haarer BK, Pringle JR. 1987. Immunofluorescence localization of the *Saccharomyces cerevisiae* CDC12 gene product to the vicinity of the 10-nm filaments in the mother-bud neck. Mol Cell Biol 7:3678–3687.

49. Halim DO, Munson M, Gao FB. 2023. The exocyst complex in neurological disorders. Hum Genet 142:1263–1270.

50. Hartwell LH. 1971. Genetic control of the cell division cycle in yeast. IV. Genes controlling bud emergence and cytokinesis. Exp Cell Res 69:265–276.

51. He B, Xi F, Zhang X, Zhang J, Guo W. 2007. Exo70 interacts with phospholipids and mediates the targeting of the exocyst to the plasma membrane. EMBO J 26:4053–4065.

52. Heider MR, Gu M, Duffy CM, Mirza AM, Marcotte LL, Walls AC, Farrall N, Hakhverdyan Z, Field MC, Rout MP, Frost A, Munson M. 2016. Subunit connectivity, assembly determinants and architecture of the yeast exocyst complex. Nat Struct Mol Biol 23:59–66.

53. Heider MR, Munson M. 2012. Exorcising the exocyst complex. Traffic 13:898–907.

54. Hernández-Rodríguez Y, Momany M. 2012. Posttranslational modifications and assembly of septin heteropolymers and higher-order structures. Curr Opin Microbiol 15:660–668.

55. Hsu SC, Hazuka CD, Roth R, Foletti DL, Heuser J, Scheller RH. 1998. Subunit composition, protein interactions, and structures of the mammalian brain sec6/8 complex and septin filaments. Neuron 20:1111–1122.

56. Hsu SC, TerBush D, Abraham M, Guo W. 2004. The exocyst complex in polarized exocytosis. Int Rev Cytol 233:243–265.

57. Hu Q, Nelson WJ. 2011. Ciliary diffusion barrier: the gatekeeper for the primary cilium compartment. Cytoskeleton (Hoboken) 68:313–324.

58. Jin Y, Sultana A, Gandhi P, Franklin E, Hamamoto S, Khan AR, Munson M, Schekman R, Weisman LS. 2011. Myosin V transports secretory vesicles via a Rab GTPase cascade and interaction with the exocyst complex. Dev Cell 21:1156–1170.

59. Jourdain I, Dooley HC, Toda T. 2012. Fission yeast Sec3 bridges the exocyst complex to the actin cytoskeleton. Traffic 13:1481–1495.

60. Jumper J, Evans R, Pritzel A, Green T, Figurnov M, Ronneberger O, Tunyasuvunakool K, Bates R, Žídek A, Potapenko A, Bridgland A, Meyer C, Kohl SAA, Ballard AJ, Cowie A, Romera-Paredes B, Nikolov S, Jain R, Adler J, Back T, et al. 2021. Highly accurate protein structure prediction with AlphaFold. Nature 596:583–589.

61. Kartmann B, Roth D. 2001. Novel roles for mammalian septins: from vesicle trafficking to oncogenesis. J Cell Sci 114:839–844.

62. Kim DU, Hayles J, Kim D, Wood V, Park HO, Won M, Yoo HS, Duhig T, Nam M, Palmer G, Han S, Jeffery L, Baek ST, Lee H, Shim YS, Lee M, Kim L, Heo KS, Noh EJ, Lee AR, et al. 2010. Analysis of a genome-wide set of gene deletions in the fission yeast *Schizosaccharomyces pombe*. Nat Biotechnol 28:617–623.

63. Kim HB, Haarer BK, Pringle JR. 1991. Cellular morphogenesis in the *Saccharomyces cerevisiae* cell cycle: localization of the CDC3 gene product and the timing of events at the budding site. J Cell Biol 112:535–544.

64. Kinoshita M. 2003. Assembly of mammalian septins. J Biochem (Tokyo) 134:491–496.

65. Kinoshita M, Kumar S, Mizoguchi A, Ide C, Kinoshita A, Haraguchi T, Hiraoka Y, Noda M. 1997. Nedd5, a mammalian septin, is a novel cytoskeletal component interacting with actin-based structures. Genes Dev 11:1535–1547.

66. Laporte D, Coffman VC, Lee I-J, Wu J-Q. 2011. Assembly and architecture of precursor nodes during fission yeast cytokinesis. J Cell Biol 192:1005–1021.

67. Lee I-J, Wu J-Q. 2012. Characterization of Mid1 domains for targeting and scaffolding in fission yeast cytokinesis. J Cell Sci 125:2973–2985.

68. Lee I-J, Wang N, Hu W, Schott K, Bähler J, Giddings TH, Jr., Pringle JR, Du L-L, Wu J-Q. 2014. Regulation of spindle pole body assembly and cytokinesis by the centrin-binding protein Sfi1 in fission yeast. Mol Biol Cell 25:2735–2749.

69. Lee PR, Song S, Ro HS, Park CJ, Lippincott J, Li R, Pringle JR, De Virgilio C, Longtine MS, Lee KS. 2002. Bni5p, a septin-interacting protein, is required for normal septin function and cytokinesis in *Saccharomyces cerevisiae*. Mol Cell Biol 22:6906–6920.

70. Lepore D, Spassibojko O, Pinto G, Collins RN. 2016. Cell cycle-dependent phosphorylation of Sec4p controls membrane deposition during cytokinesis. J Cell Biol 214:691–703.

71. Lepore DM, Martínez-Núñez L, Munson M. 2018. Exposing the elusive exocyst structure. Trends Biochem Sci 43:714–725.

72. Li CR, Lee RT, Wang YM, Zheng XD, Wang Y. 2007. *Candida albicans* hyphal morphogenesis occurs in Sec3p-independent and Sec3p-dependent phases separated by septin ring formation. J Cell Sci 120:1898–1907.

73. Lippincott J, Li R. 1998. Sequential assembly of myosin II, an IQGAP-like protein, and filamentous actin to a ring structure involved in budding yeast cytokinesis. J Cell Biol 140:355–366.

74. Liu D, Li X, Shen D, Novick P. 2018. Two subunits of the exocyst, Sec3p and Exo70p, can function exclusively on the plasma membrane. Mol Biol Cell 29:736–750.

75. Liu J, Guo W. 2012. The exocyst complex in exocytosis and cell migration. Protoplasma 249:587–597.

76. Liu J, Wang H, McCollum D, Balasubramanian MK. 1999. Drc1p/Cps1p, a 1,3-β-glucan synthase subunit, is essential for division septum assembly in *Schizosaccharomyces pombe*. Genetics 153:1193–1203.

77. Longtine MS, Bi E. 2003. Regulation of septin organization and function in yeast. Trends Cell Biol 13:403–409.

78. Longtine MS, DeMarini DJ, Valencik ML, Al-Awar OS, Fares H, De Virgilio C, Pringle JR. 1996. The septins: roles in cytokinesis and other processes. Curr. Opin. Cell Biol. 8:106–119.

79. Longtine MS, Fares H, Pringle JR. 1998. Role of the yeast Gin4p protein kinase in septin assembly and the relationship between septin assembly and septin function. J Cell Biol 143:719–736.

80. Luo G, Zhang J, Guo W. 2014. The role of Sec3p in secretory vesicle targeting and exocyst complex assembly. Mol Biol Cell 25:3813–3822.

81. Marquardt J, Chen X, Bi E. 2019. Architecture, remodeling, and functions of the septin cytoskeleton. Cytoskeleton (Hoboken) 76:7–14.

82. Martin-Cuadrado AB, Duenas E, Sipiczki M, Vazquez de Aldana CR, del Rey F. 2003. The endo-β-1,3-glucanase eng1p is required for dissolution of the primary septum during cell separation in *Schizosaccharomyces pombe*. J Cell Sci 116:1689–1698.

83. Martin-Cuadrado AB, Morrell JL, Konomi M, An H, Petit C, Osumi M, Balasubramanian M, Gould KL, Del Rey F, de Aldana CR. 2005. Role of septins and the exocyst complex in the function of hydrolytic enzymes responsible for fission yeast cell separation. Mol Biol Cell 16:4867–4881.

84. Martin-Urdiroz M, Deeks MJ, Horton CG, Dawe HR, Jourdain I. 2016. The exocyst complex in health and disease. Front Cell Dev Biol 4:24.

85. McMurray MA. 2019. The long and short of membrane curvature sensing by septins. J Cell Biol 218:1083–1085.

86. McMurray MA, Bertin A, Garcia G, III, Lam L, Nogales E, Thorner J. 2011. Septin filament formation is essential in budding yeast. Dev Cell 20:540–549.

87. McMurray MA, Thorner J. 2019. Turning it inside out: The organization of human septin heterooligomers. Cytoskeleton (Hoboken). 76:449–456.

88. Mei K, Guo W. 2018. The exocyst complex. Curr Biol 28:R922–R925.

89. Mei K, Li Y, Wang S, Shao G, Wang J, Ding Y, Luo G, Yue P, Liu J-J, Wang X, Dong M-Q, Wang H-W, Guo W. 2018. Cryo-EM structure of the exocyst complex. Nat Struct Mol Biol 25:139–146.

90. Meitinger F, Palani S. 2016. Actomyosin ring driven cytokinesis in budding yeast. Semin Cell Dev Biol 53:19–27.

91. Meitinger F, Palani S, Hub B, Pereira G. 2013. Dual function of the NDR-kinase Dbf2 in the regulation of the F-BAR protein Hof1 during cytokinesis. Mol Biol Cell 24:1290–1304.

92. Mendonça DC, Macedo JN, Guimarães SL, Barroso da Silva FL, Cassago A, Garratt RC, Portugal RV, Araujo APU. 2019. A revised order of subunits in mammalian septin complexes. Cytoskeleton (Hoboken). 76:457–466.

93. Michaelis AC, Brunner AD, Zwiebel M, Meier F, Strauss MT, Bludau I, Mann M. 2023. The social and structural architecture of the yeast protein interactome. Nature 624:192–200.

94. Mirdita M, Schütze K, Moriwaki Y, Heo L, Ovchinnikov S, Steinegger M. 2022. ColabFold: making protein folding accessible to all. Nat Methods 19:679–682.

95. Momany M, Talbot NJ. 2017. Septins focus cellular growth for host infection by pathogenic fungi. Front Cell Dev Biol 5:33.

96. Moreno S, Klar A, Nurse P. 1991. Molecular genetic analysis of fission yeast *Schizosaccharomyces pombe*. Methods Enzymol 194:795–823.

97. Mostowy S, Cossart P. 2012. Septins: the fourth component of the cytoskeleton. Nat Rev Mol Cell Biol 13:183–194.

98. Muñoz S, Manjón E, Sánchez Y. 2014. The putative exchange factor Gef3p interacts with Rho3p GTPase and the septin ring during cytokinesis in fission yeast. J Biol Chem 289:21995–22007.

99. Neubauer K, Zieger B. 2021. Role of septins in endothelial cells and platelets. Front Cell Dev Biol 9:768409.

100. Neufeld TP, Rubin GM. 1994. The Drosophila *peanut* gene is required for cytokinesis and encodes a protein similar to yeast putative bud neck filament proteins. Cell 77:371–379.

101. Nishihama R, Onishi M, Pringle JR. 2011. New insights into the phylogenetic distribution and evolutionary origins of the septins. Biol Chem 392:681–687.

102. Oh Y, Bi E. 2011. Septin structure and function in yeast and beyond. Trends Cell Biol 21:141–148.

103. Oh Y, Schreiter J, Nishihama R, Wloka C, Bi E. 2013. Targeting and functional mechanisms of the cytokinesis-related F-BAR protein Hof1 during the cell cycle. Mol Biol Cell 24:1305–1320.

104. Okada S, Leda M, Hanna J, Savage NS, Bi E, Goryachev AB. 2013. Daughter cell identity emerges from the interplay of Cdc42, septins, and exocytosis. Dev Cell 26:148–161.

105. Onishi M, Koga T, Hirata A, Nakamura T, Asakawa H, Shimoda C, Bähler J, Wu J-Q, Takegawa K, Tachikawa H, Pringle JR, Fukui Y. 2010. Role of septins in the orientation of forespore membrane extension during sporulation in fission yeast. Mol Cell Biol 30:2057–2074.

106. Onishi M, Pringle JR. 2016. The nonopisthokont septins: How many there are, how little we know about them, and how we might learn more. Methods Cell Biol 136:1–19.

107. Paiano A, Margiotta A, De Luca M, Bucci C. 2019. Yeast two-hybrid assay to identify interacting proteins. Curr Protoco Protein Sci 95:e70.

108. Perez AM, Finnigan GC, Roelants FM, Thorner J. 2016. Septin-associated protein kinases in the yeast *Saccharomyces cerevisiae*. Front Cell Dev Biol 4:119.

109. Perez P, Portales E, Santos B. 2015. Rho4 interaction with exocyst and septins regulates cell separation in fission yeast. Microbiology 161:948–959.

110. Petit CS, Mehta S, Roberts RH, Gould KL. 2005. Ace2p contributes to fission yeast septin ring assembly by regulating *mid2^+^* expression. J Cell Sci 118:5731–5742.

111. Ren J, Guo W. 2012. ERK1/2 regulate exocytosis through direct phosphorylation of the exocyst component Exo70. Dev Cell 22:967–978.

112. Russell SE, Hall PA. 2005. Do septins have a role in cancer? Br J Cancer 93:499–503.

113. Safavian D, Kim MS, Xie H, El-Zeiry M, Palander O, Dai L, Collins RF, Froese C, Shannon R, Nagata KI, Trimble WS. 2023. Septin-mediated RhoA activation engages the exocyst complex to recruit the cilium transition zone. J Cell Biol 222:e201911062.

114. Santos B, Martin-Cuadrado AB, Vazquez de Aldana CR, del Rey F, Perez P. 2005. Rho4 GTPase is involved in secretion of glucanases during fission yeast cytokinesis. Eukaryot Cell 4:1639–1645.

115. Sharma K, Menon MB. 2023. Decoding post-translational modifications of mammalian septins. Cytoskeleton (Hoboken) 80:169–181.

116. Sheffield PJ, Oliver CJ, Kremer BE, Sheng S, Shao Z, Macara IG. 2003. Borg/septin interactions and the assembly of mammalian septin heterodimers, trimers, and filaments. J Biol Chem 278:3483–3488.

117. Shen D, Yuan H, Hutagalung A, Verma A, Kümmel D, Wu X, Reinisch K, McNew JA, Novick P. 2013. The synaptobrevin homologue Snc2p recruits the exocyst to secretory vesicles by binding to Sec6p. J Cell Biol 202:509–526.

118. Shi W, Cannon KS, Curtis BN, Edelmaier C, Gladfelter AS, Nazockdast E. 2023. Curvature sensing as an emergent property of multiscale assembly of septins. Proc Natl Acad Sci U S A 120:e2208253120.

119. Sivaram MV, Saporita JA, Furgason ML, Boettcher AJ, Munson M. 2005. Dimerization of the exocyst protein Sec6p and its interaction with the t-SNARE Sec9p. Biochemistry 44:6302–6311.

120. Sjölinder M, Uhlmann J, Ponstingl H. 2002. DelGEF, a homologue of the Ran guanine nucleotide exchange factor RanGEF, binds to the exocyst component Sec5 and modulates secretion. FEBS Lett 532:211–215.

121. Soroor F, Kim MS, Palander O, Balachandran Y, Collins RF, Benlekbir S, Rubinstein JL, Trimble WS. 2021. Revised subunit order of mammalian septin complexes explains their in vitro polymerization properties. Mol Biol Cell. 32:289–300.

122. Synek L, Pleskot R, Sekereš J, Serrano N, Vukašinović N, Ortmannová J, Klejchová M, Pejchar P, Batystová K, Gutkowska M, Janková-Drdová E, Marković V, Pečenková T, Šantrůček J, Žárský V, Potocký M. 2021. Plasma membrane phospholipid signature recruits the plant exocyst complex via the EXO70A1 subunit. Proc Natl Acad Sci U S A 118:e2105287118.

123. Tasto JJ, Morrell JL, Gould KL. 2003. An anillin homologue, Mid2p, acts during fission yeast cytokinesis to organize the septin ring and promote cell separation. J Cell Biol 160:1093–1103.

124. Tay YD, Leda M, Spanos C, Rappsilber J, Goryachev AB, Sawin KE. 2019. Fission yeast NDR/LATS kinase Orb6 regulates exocytosis via phosphorylation of the exocyst complex. Cell Rep 26:1654–1667.

125. TerBush DR, Maurice T, Roth D, Novick P. 1996. The Exocyst is a multiprotein complex required for exocytosis in *Saccharomyces cerevisiae*. EMBO J 15:6483–6494.

126. TerBush DR, Novick P. 1995. Sec6, Sec8, and Sec15 are components of a multisubunit complex which localizes to small bud tips in *Saccharomyces cerevisiae*. J Cell Biol 130:299–312.

127. Tokhtaeva E, Capri J, Marcus EA, Whitelegge JP, Khuzakhmetova V, Bukharaeva E, Deiss-Yehiely N, Dada LA, Sachs G, Fernandez-Salas E, Vagin O. 2015. Septin dynamics are essential for exocytosis. J Biol Chem 290:5280–5297.

128. Vallen EA, Caviston J, Bi E. 2000. Roles of Hof1p, Bni1p, Bnr1p, and Myo1p in cytokinesis in *Saccharomyces cerevisiae*. Mol Biol Cell 11:593–611.

129. Vavylonis D, Wu J-Q, Hao S, O’Shaughnessy B, Pollard TD. 2008. Assembly mechanism of the contractile ring for cytokinesis by fission yeast. Science 319:97–100.

130. Vega IE, Hsu SC. 2003. The septin protein Nedd5 associates with both the exocyst complex and microtubules and disruption of its GTPase activity promotes aberrant neurite sprouting in PC12 cells. Neuroreport 14:31–37.

131. Versele M, Thorner J. 2005. Some assembly required: yeast septins provide the instruction manual. Trends Cell Biol 15:414–424.

132. Wang H, Tang X, Balasubramanian MK. 2003. Rho3p regulates cell separation by modulating exocyst function in *Schizosaccharomyces pombe*. Genetics 164:1323–1331.

133. Wang H, Tang X, Liu J, Trautmann S, Balasundaram D, McCollum D, Balasubramanian MK. 2002. The multiprotein exocyst complex is essential for cell separation in *Schizosaccharomyces pombe*. Mol Biol Cell 13:515–529.

134. Wang N, Lee I-J, Rask G, Wu J-Q. 2016. Roles of the TRAPP-II complex and the exocyst in membrane deposition during fission yeast cytokinesis. PLoS Biol 14:e1002437.

135. Wang N, Lo Presti L, Zhu Y-H, Kang M, Wu Z, Martin SG, Wu J-Q. 2014. The novel proteins Rng8 and Rng9 regulate the myosin-V Myo51 during fission yeast cytokinesis. J Cell Biol 205:357–375.

136. Wang N, Wang M, Zhu Y-H, Grosel TW, Sun D, Kudryashov DS, Wu J-Q. 2015. The Rho-GEF Gef3 interacts with the septin complex and activates the GTPase Rho4 during fission yeast cytokinesis. Mol Biol Cell 26:238–255.

137. Werner B, Yadav S. 2023. Phosphoregulation of the septin cytoskeleton in neuronal development and disease. Cytoskeleton (Hoboken) 80:275–289.

138. Wloka C, Nishihama R, Onishi M, Oh Y, Hanna J, Pringle JR, Krauss M, Bi E. 2011. Evidence that a septin diffusion barrier is dispensable for cytokinesis in budding yeast. Biol Chem 392:813–829.

139. Woods A, Sherwin T, Sasse R, MacRae TH, Baines AJ, Gull K. 1989. Definition of individual components within the cytoskeleton of *Trypanosoma brucei* by a library of monoclonal antibodies. J Cell Sci 93:491–500.

140. Woods BL, Gladfelter AS. 2021. The state of the septin cytoskeleton from assembly to function. Curr Opin Cell Biol 68:105–112.

141. Wu J-Q, Kuhn JR, Kovar DR, Pollard TD. 2003. Spatial and temporal pathway for assembly and constriction of the contractile ring in fission yeast cytokinesis. Dev Cell 5:723–734.

142. Wu J-Q, Ye Y, Wang N, Pollard TD, Pringle JR. 2010. Cooperation between the septins and the actomyosin ring and role of a cell-integrity pathway during cell division in fission yeast. Genetics 186:897–915.

143. Yang L, Zhang X, Gu Y, Shi Y, Wang LB, Shi JX, Zhen XX, Xin YW, Gu WW, Wang J. 2022. SEC5 is involved in M2 polarization of macrophages via the STAT6 pathway, and its dysfunction in decidual macrophages is associated with recurrent spontaneous abortion. Front Cell Dev Biol 10:891748.

144. Ye Y, Lee I-J, Runge KW, Wu J-Q. 2012. Roles of putative Rho-GEF Gef2 in division-site positioning and contractile-ring function in fission yeast cytokinesis. Mol Biol Cell 23:1181–1195.

145. Ye Y, Osmani AH, Liu Z-R, Kern A, Wu J-Q. 2025. Fission yeast GPI inositol deacylase Bst1 regulates ER-Golgi transport and functions in late stages of cytokinesis. Mol Biol Cell 36:ar27.

146. Yin R, Pierce BG. 2024. Evaluation of AlphaFold antibody-antigen modeling with implications for improving predictive accuracy. Protein Sci. 33:e4865.

147. Yue P, Zhang Y, Mei K, Wang S, Lesigang J, Zhu Y, Dong G, Guo W. 2017. Sec3 promotes the initial binary t-SNARE complex assembly and membrane fusion. Nat Commun 8:14236.

148. Zhang L, Cai Y, Li Y, Zhang T, Wang B, Lu G, Zhang D, Olsson S, Wang Z. 2021. MoSep3 and MoExo70 are needed for MoCK2 ring assembly essential for appressorium function in the rice blast fungus, *Magnaporthe oryzae*. Mol Plant Pathol 22:1159–1164.

149. Zhang XM, Ellis S, Sriratana A, Mitchell CA, Rowe T. 2004. Sec15 is an effector for the Rab11 GTPase in mammalian cells. J Biol Chem 279:43027–43034.

150. Zheng S, Dong F, Rasul F, Yao X, Jin QW, Zheng F, Fu C. 2018. Septins regulate the equatorial dynamics of the separation initiation network kinase Sid2p and glucan synthases to ensure proper cytokinesis. FEBS J 285:2468–2480.

151. Zheng S, Zheng B, Fu C. 2024. The roles of septins in regulating fission yeast cytokinesis. J Fungi 10:115.

152. Zhou TT, Zhao YL, Guo HS. 2017. Secretory proteins are delivered to the septin-organized penetration interface during root infection by *Verticillium dahliae*. PLoS Pathog 13:e1006275.

153. Zhu Y-H, Hyun J, Pan YZ, Hopper JE, Rizo J, Wu J-Q. 2018. Roles of the fission yeast UNC-13/Munc13 protein Ync13 in late stages of cytokinesis. Mol Biol Cell 29:2259–2279.

154. Zhu Y-H, Ye Y, Wu Z, Wu J-Q. 2013. Cooperation between Rho-GEF Gef2 and its binding partner Nod1 in the regulation of fission yeast cytokinesis. Mol Biol Cell 24:3187–3204.

